# Expression of the MSPDBL2 antigen in a discrete subset of *Plasmodium falciparum* schizonts is regulated by GDV1 but not linked to sexual commitment

**DOI:** 10.1101/2023.11.21.568010

**Authors:** Aline Freville, Lindsay B. Stewart, Kevin K.A. Tetteh, Moritz Treeck, Alfred Cortes, Till S. Voss, Sarah J. Tarr, David A. Baker, David J. Conway

## Abstract

The *Plasmodium falciparum* merozoite surface protein MSPDBL2 is a polymorphic antigen targeted by acquired immune responses, and normally expressed in only a minority of mature schizonts. The potential relationship of MSPDBL2 to sexual commitment is examined, as variable *mspdbl2* transcript levels and proportions of MSPDBL2-positive mature schizonts in clinical isolates have previously correlated with levels of many sexual stage parasite gene transcripts, although not with the master regulator *ap2-g*. It is demonstrated that conditional overexpression of GDV1, which promotes sexual commitment, also substantially increases the proportion of MSPDBL2-positive schizonts in culture. Conversely, truncation of the *gdv1* gene is shown to prevent any expression of MSPDBL2. However, across diverse *P. falciparum* cultured lines the variable proportions of MSPDBL2 positivity in schizonts does not correlate significantly with variable gametocyte conversion rates, indicating it is not involved in sexual commitment. Confirming this, examining a line with endogenous HA-tagged AP2-G showed that the individual schizonts expressing MSPDBL2 are mostly different to those expressing AP2-G. Using a selection-linked integration system, modified *P. falciparum* lines were engineered to express an intact or disrupted version of MSPDBL2, showing the protein is not required for sexual commitment or early gametocyte development. Asexual parasite multiplication rates were also not affected by expression of either intact or disrupted MSPDBL2 in a majority of schizonts. Occurring alongside sexual commitment, the role of the discrete MSPDBL2-positive schizont subpopulation requires further investigation in natural infections where it is under immune selection.

## INTRODUCTION

Asexual blood stage malaria parasites are under selection from acquired immune responses, and clonally vary expression of some target antigens on the infected erythrocyte surface and on the merozoite that invades erythrocytes (1, 2). The *Plasmodium falciparum* merozoite surface protein MSPDBL2 is a target of naturally acquired antibodies against conserved as well as allele-specific epitopes (3, 4) and subject to strong balancing selection within endemic populations (5, 6). Two population cohort studies have indicated that MSPDBL2 antibodies in plasma of children in endemic populations were associated with reduced prospective risk of developing clinical malaria (3, 7). MSPDBL2 is a member of the MSP3-like protein family, which is defined by the presence of a conserved N-terminal five amino acid motif (8) and a C-terminal domain believed to be involved in protein oligomerization (9, 10). MSPDBL2 and MSPDBL1 are the only MSP3-like members to possess a cysteine-rich Duffy-binding-like (DBL) domain (7, 8) which enables these proteins to interact with erythrocytes (11), although no specific host receptor has been identified so far (11, 12).

During *P. falciparum* schizont development, the MSPDBL2 protein is expressed and released into the parasitophorous vacuole, becoming associated with the merozoite surface through interaction with MSP1 (11, 12). Although MSPDBL2 on the merozoite surface has been suggested to be involved in erythrocyte invasion (11), disruption of the *mspdbl2* gene previously has shown no effect on asexual replication in culture (13). Pertinent to this observation, the *mspdbl2* gene is epigenetically regulated and usually in a suppressed state, carrying a H3K9me3/heterochromatin protein 1 (HP1)-marked signature of heterochromatin (5, 14, 15). Analysis of schizont stage cultures of clinical isolates and long-term *in vitro* culture-adapted lines have shown that *mspdbl2* transcript levels vary substantially among isolates (5, 16). Moreover, the MSPDBL2 protein is only expressed in a subpopulation of mature schizonts within culture-adapted parasite clones (5) as well as in first cycle *ex vivo* culture of clinical isolates (17).

The DBL domain of the MSPDBL2 protein has been shown to directly interact with a conserved region in human IgM antibodies, an interaction that might inhibit specific binding of immune IgG to MSPDBL2 during infection (18). Allelic variation in MSPDBL2 did not affect this binding (18), but *mspdbl2* gene variants have been reported to be associated with parasite sensitivity to *in vitro* inhibition by halofantrine, mefloquine and lumefantrine (19–21). It has been reported that the over-expression of GDV1, a nuclear protein activating expression of the master regulator of sexual commitment AP2-G, also rapidly increases the transcription of a small set of other genes including *mspdbl2* (22). Although it was not shown that GDV1 directly affects MSPDBL2 protein expression, this observation has led to a hypothesis that MSPDBL2 may be specifically expressed in sexually committed parasites (16, 22).

Here, it is shown that GDV1 overexpression substantially increases not only the rate of sexual conversion but also the proportion of schizonts expressing the MSPDBL2 protein. Furthermore, expression of GDV1 is essential for the normal expression of MSPDBL2 in a minority of schizonts, as truncation of the *gdv1* gene ablated this. However, an extensive analysis across diverse *P. falciparum* cultured lines showed no correlation between variable proportions of schizonts expressing MSPDBL2 and the rates of sexual conversion in culture. Furthermore, immunofluorescence microscopy showed that the normal minority proportions of individual schizonts which express either MSPDBL2 or AP2-G are mostly different, indicating that MSPDBL2 is not a marker of sexual commitment. Selection-linked integration was used to engineer parasite cultures to express either an intact or a truncated version of MSPDBL2 in most schizonts, which confirmed that sexual commitment and early gametocyte development do not require intact MSPDBL2, while asexual multiplication rates were also unaffected. Thus, GDV1 is involved in the regulation of a discrete antigenic subpopulation that is mostly asexually committed and under strong immune selection in natural infections, which warrants further investigation in clinical and population-based studies.

## RESULTS

### GDV1 expression significantly increases the proportion of MSPDBL2-positive schizonts

To test the effect of GDV1 overexpression on the proportion of mature schizonts expressing MSPDBL2, we used the genetically engineered *P. falciparum* 3D7/GDV1-GFP-DD parasite line (3D7/iGP). In this line, GDV1 is over-expressed from an ectopic locus when parasites are cultured in the presence of the stabilising reagent Shield-1 (23). As expected, in four independent experiments, live fluorescence microscopy showed that most trophozoites (a mean of 76.2%) expressed GDV1 in the presence of 1 µM Shield-1, compared to only 2.9% in the absence of Shield-1 (Figure 1A). In each of these experiments, the proportion of schizonts that were MSPDBL2-positive (MSPDBL2+) was also massively increased by GDV1 over-expression, the mean proportion being 38.5% with Shield-1, and only 1.1% without Shield-1 (Figure 1A, Supplementary Table S1). Additional tests were made using exposure to intermediate concentrations of Shield-1 reagent, which yielded a range of intermediate proportions of MSPDBL2+ schizonts (Figure 1B, Supplementary Table S2). The gametocyte conversion rates of these cultures had previously been assayed by examining parasites in the next cycle using Pfs16 staining (published data (24) tabulated here in Supplementary Table S2). There was a significantly positive correlation between these gametocyte conversion rates and the proportions of MSPDBL2+ schizonts (Spearman’s rho = 0.65, P = 0.0095)(Figure 1C), showing that both responded in parallel to the overexpression of GDV1.

**Figure 1.**
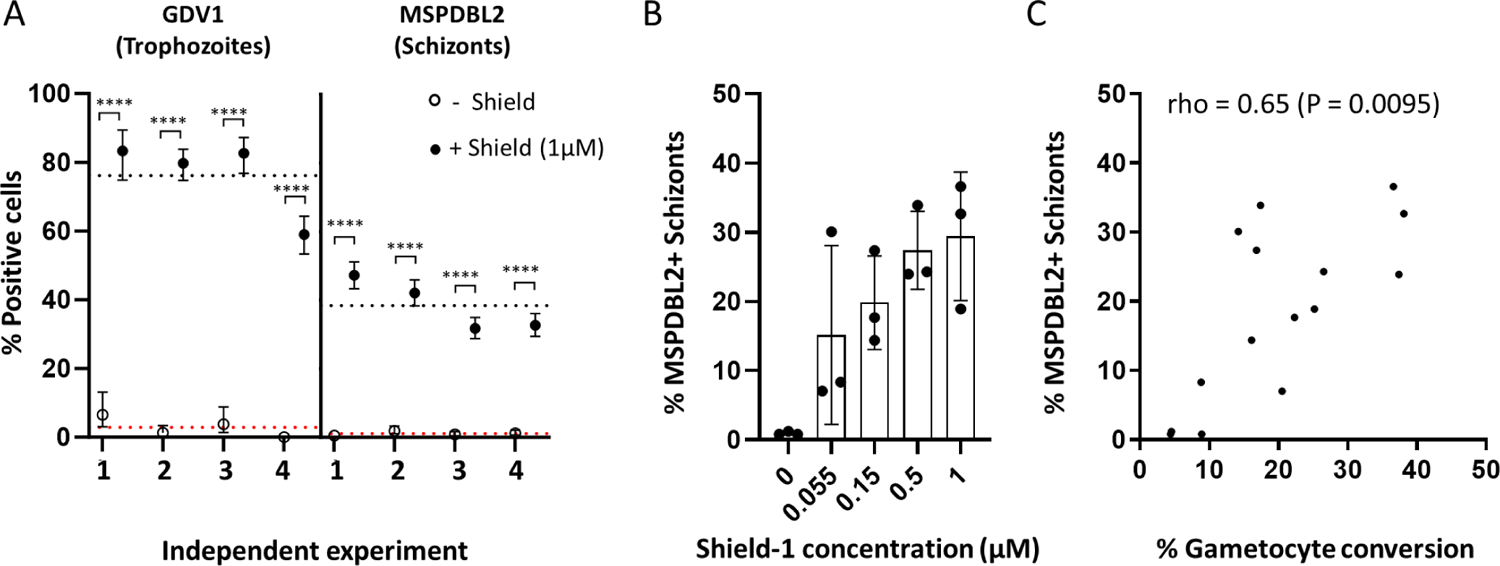
GDV1 overexpression increases the proportion of MSPDBL2+ schizonts. **A.** Proportions of trophozoites expressing GDV1 and mature schizonts expressing MSPDBL2 (with 95% confidence intervals) in the 3D7/iGP parasite line cultured in presence (closed circles) or absence (open circles) of 1µM Shield-1 reagent. Four independent experimental replicates were performed. Asterisks indicate significant differences between paired measurements in each of seven biological replicate assays (Fisher’s Exact tests, P ˂ 0.0001). Dotted lines represent the mean across replicate assays (red, no Shield; black, + 1µM Shield). All individual counts and assay values are given in Supplementary Table S1. **B.** Varying proportions of MSPDBL2+ schizonts in 3D7/iGP parasites cultured with different concentrations of Shield, bars showing means with standard deviation of three biological replicate assays performed for each concentration. All individual counts and assay values are given in Supplementary Table S2. **C.** Significant correlation between MSPDBL2+ schizont proportions and the gametocyte conversion rate across all assays with various concentrations of Shield (n= 15, details in Supplementary Table S2).

### Truncation of the *gdv1* gene prevents MSPDBL2 expression in schizonts

MSPDBL2 is normally only expressed in a small minority of schizonts in *P. falciparum*. To test whether GDV1 is required for this normal expression, parasites in which the *gdv1* gene was disrupted were investigated. The *P. falciparum* clone NF54-GDV1delta39 contains a premature stop codon that results in a truncated version of GDV1 which is inactive and prevents induction of sexual commitment (25). In three separate replicate experiments, the NF54-GDV1delta39 clone was compared with the parental NF54 clone. No MSPDBL2+ schizonts were seen in the NF54-GDV1delta39 line in any of the replicates, whereas the NF54 control parasites showed the typical minority proportion of positive mature schizonts (between 0.6 and 1.3% for each replicate culture) (Figure 2, Supplementary Table S3).

**Figure 2.**
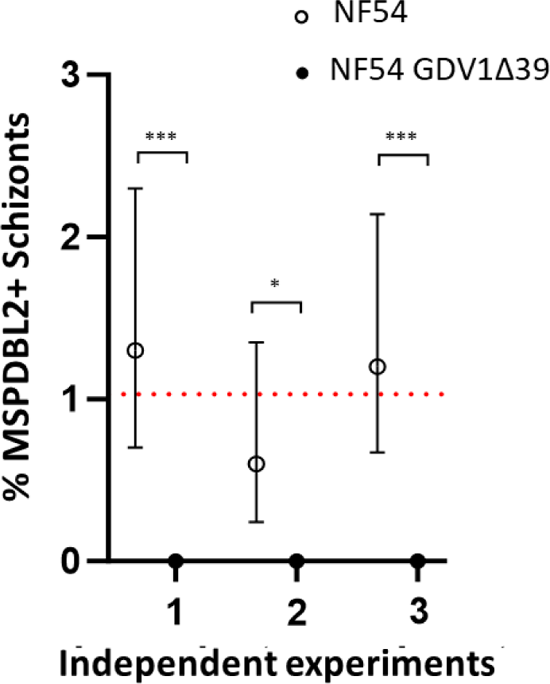
Normal MSPDBL2 expression in schizonts requires intact *gdv1*. Proportions of mature schizonts expressing MSPDBL2 (with 95% confidence intervals) in the NF54-GDV1Δ39 clone compared with the parental NF54 clone. Asterisks indicate significant differences between paired measurements in each of three biological replicate assays (Fisher’s Exact tests, *: P ˂ 0.05, ***: P ˂ 0.001). Dotted lines represent the mean across replicate assays (NF54: 1.0, NF54-GDV1Δ39:0). All individual counts and assay values are given in Supplementary Table S3.

### Varying proportions of MSPDBL2-positive schizonts among diverse *P. falciparum* laboratory lines

It was previously shown that the proportion of mature schizonts which express MSPDBL2 varies among *P. falciparum* lines (5). To gain a better view of MSPDBL2 variation in culture, multiple biological replicate preparations of each of 13 *P. falciparum* long-term culture adapted lines were analysed (Figure 3A), all lines being grown in identical serum-free medium supplemented with Albumax II. Each of these replicates was sampled from a culture in which the gametocyte conversion rate in the subsequent cycle was measured and previously reported (24), with the slides of schizonts having been archived for immunofluorescence analysis of MSPDBL2 expression in the current study.

**Figure 3.**
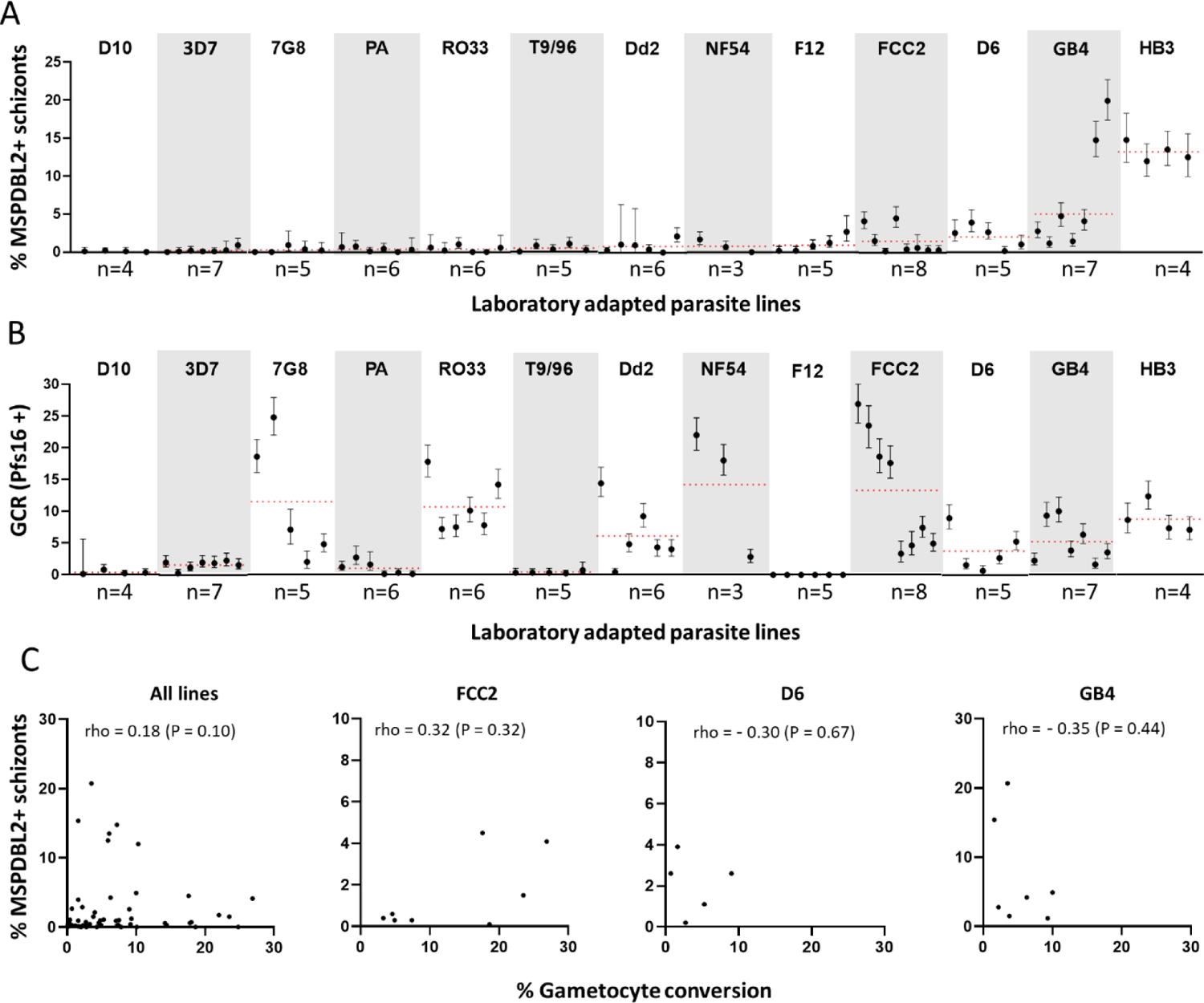
Variable proportions of schizonts expressing MSPDBL2 in different *P. falciparum* lines. **A.** For each of 13 long-term culture-adapted lines, the proportions (with 95% confidence intervals) are shown for multiple biological replicate cultures (between three and eight of each). Red dotted lines represent the mean across replicates for each parasite line. All individual counts and assay values are given in Supplementary Table S3. **B.** Gametocyte conversion rates (GCR) in the same replicate cultures, measured as the proportion of stage 1 gametocytes developing in the following cycle as determined by Pfs16 staining (GCR data previously reported (24)). **C.** No significant correlation (estimated by Spearman’s rank correlation) between MSPDBL2+ schizont proportions and the gametocyte conversion rate, across all replicates of the different *P. falciparum* lines (n= 71 paired assays across all lines), or for individual lines that showed most variation in proportions (n= 8 for FCC2, n=5 for D6, n=8 for GB4) (all numbers are given in Supplementary Table S4).

Most *P. falciparum* lines (D10, 3D7, NF54, 7G8, Palo Alto, RO33, T996, Dd2, and F12) consistently showed very low proportions of MSPDBL2+ schizonts (< 1% positive as means in immunofluorescence using antibodies specific for a conserved region of MSPDBL2) (Figure 3A, Supplementary Table S4). It is notable that no parasite line expressed zero MSPDBL2+ schizonts, and that even a parasite line containing an *ap2-g* nonsense mutation (clone F12, previously derived from 3D7) expressed MSPDBL2 in 0.9% of schizonts. This indicates that MSPDBL2 expression is not dependent on AP2-G. Two of the parasite lines showed slightly higher proportions of MSPDBL2+ schizonts (means of 1.5% in FCC2 and 2.1% in D6), and two lines showed much higher proportions (means of 7.3% in GB4 and 13.2% in HB3) (Figure 3A, Supplementary Table S4), all of these being significantly higher than in those listed above (pairwise Mann-Whitney tests, Supplementary Table S5).

The gametocyte conversion rates were previously assayed in each of these individual replicate cultures, using Pfs16 staining to discriminate early gametocytes in the following cycle (24), so these were compared with the proportions of MSPDBL2+ schizonts. All of the parasite lines except F12 produce some early gametocytes, although some of the lines produce very few (Figure 3B, Supplementary Table S4). Notably, proportions of MSPDBL2+ schizonts were not significantly correlated with the gametocyte conversion rates. For example, there are very low proportions of MSPDBL2+ schizonts in several of the parasite lines that show relatively high rates of gametocyte conversion (lines 7G8, RO33, Dd2, NF54 and FCC2) (Figure 3A and 3B). In a pooled analysis across all parasite lines, there was only a very weak positive correlation between proportions of MSPDBL2+ schizonts and gametocyte conversion rates (Spearman’s rho = 0.18, P = 0.10, Figure 3C). Considering inter-replicate variation in rates for individual lines, separate analyses of each of the three parasite lines with most inter-replicate variation in proportions of MSPDBL2+ schizonts (FCC2, D6, and GB4) showed no significant correlations with the gametocyte conversion rates of the same culture replicates (Figure 3C, Supplementary Table S4). As routine estimates of parasitaemia by Giemsa staining of slides in culture maintenance had been recorded for 42 of these cultures in both cycles (Supplementary Table S4), the possibility of correlations between proportions of MSPDBL2+ schizonts and the parasitaemia levels or fold-changes between cycles was also explored. This revealed no significant correlation, and no directional trend for variation within each of the parasite lines that had at least five replicates (mean rho = 0.21, P values > 0.56 for MSPDBL2 expression in schizonts versus parasitaemia in the same cycle; mean rho = −0.09, all P values > 0.26 for MSPDBL2 expression in schizonts and parasitaemia fold change) (Supplementary Table S6).

### Effect of choline on MSPDBL2-positive schizont proportions

Choline supplementation of serum-free medium has been previously shown to suppress parasite GDV1 expression (22) and sexual commitment rates (22, 24, 26, 27). Given that GDV1 expression increases MSPDBL2+ schizont proportions in the inducible parasite line 3D7/iGP, it is relevant to test for a potentially suppressive effect of choline supplementation on proportions of MSPDBL2+ schizonts. The HB3 and NF54 lines were analysed here, as they have contrasting proportions of MSPDBL2+ schizonts (relatively high in HB3 and very low in NF54) (Figure 3). Multiple biological assay replicates were performed on each of these lines to test the effect of culture medium to which either 2 mM choline or no choline was added, with parasites exposed from the ring stage of development onwards and allowed to develop to schizonts for analysis of MSPDBL2 expression.

The addition of 2 mM choline significantly suppressed the proportion of MSPDBL2+ schizonts in four of seven replicates performed on the HB3 parasite line (P < 0.001 for each of the four replicates, Figure 4A, Supplementary Table S7). All four replicates with significant effects had >10% MSPDBL2+ schizonts in the controls without choline treatment, whereas the three replicates that showed no effect had significantly lower MSPDBL2+ schizont proportions in the controls (Fisher’s Exact P = 0.029 in test for association between initial proportions positive and effect of choline). For NF54, which had very low MSPDBL2+ schizont proportions in all controls, there was no significant effect of choline in reducing these proportions in any of the six replicate assays performed. These results indicate that adding choline suppresses proportions of MSPDBL2+ schizonts under conditions in which these would otherwise be high, consistent with the previously known effect of choline suppressing GDV1 expression. The effect of choline in suppressing gametocyte conversion rates was previously reported for these same culture replicates (24), showing significant effects in most replicates of both parasite lines (Figure 4B).

**Figure 4.**
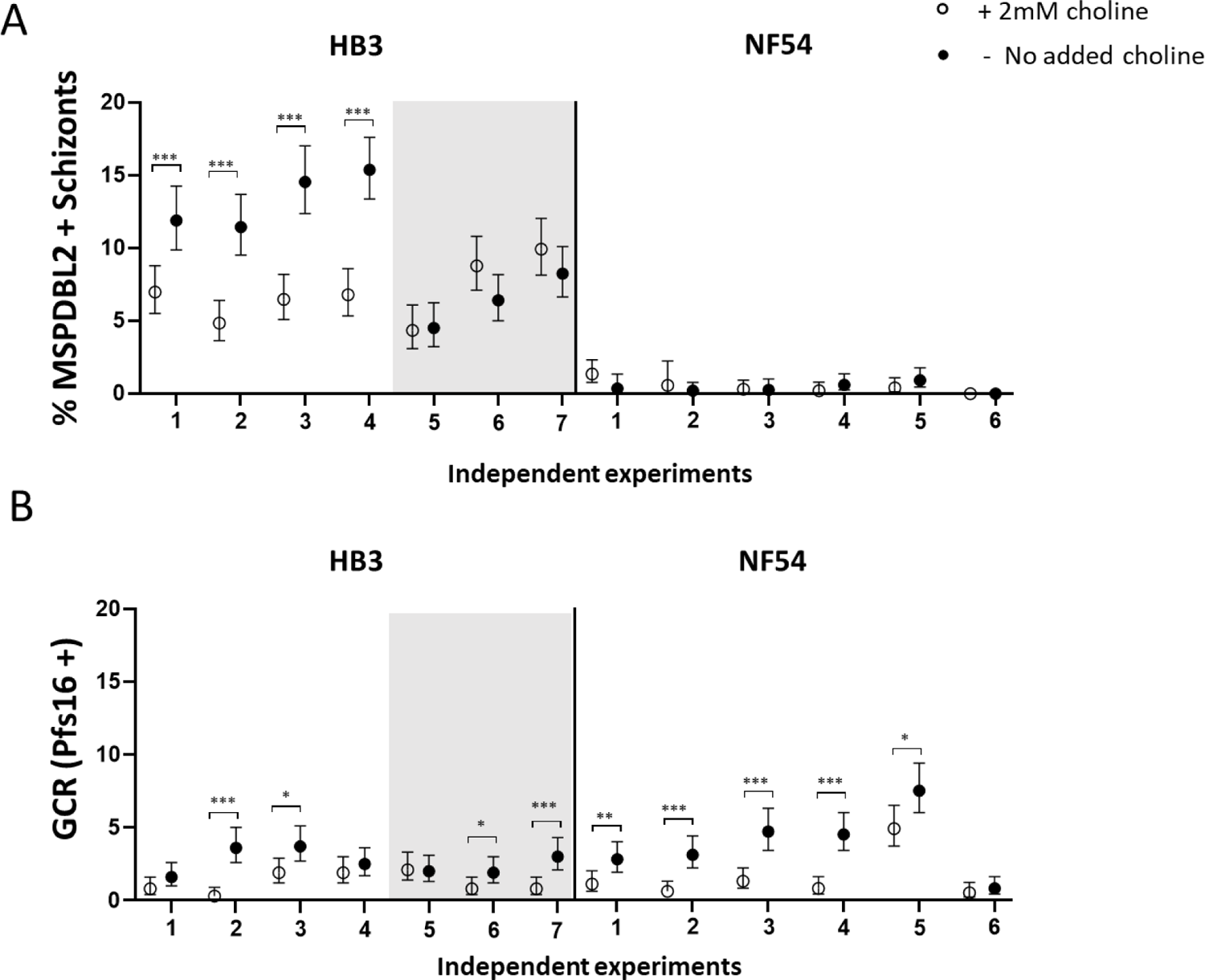
Effect of choline on proportions of mature schizonts expressing MSPDBL2. **A.** Multiple independent assays were performed on two *P. falciparum lines* with contrasting proportions of MSPDBL2+ schizonts. For line HB3, adding choline to serum-free medium reduced the proportions of schizonts expressing MSLDBL2 in four of the experimental replicate experiments, in each of which the proportions in controls were >10%. The other three replicates of HB3 (shown against shaded background) had lower proportions in controls and choline did not reduce these. In parasite line NF54, proportions of schizonts expressing MSPDBL2 were consistently very low, with or without added choline. (Fisher’s Exact Tests * P < 0.05, * *P < 0.01, *** P < 0.001). **B.** Gametocyte conversion rates (GCR) measured in the following cycle of the same replicate cultures using anti-Pfs16 staining (GCR data previously reported (24)). Choline showed suppressive effects on GCR in most experimental replicates in both lines. All individual counts and assay values for all biological replicate assay are given in Supplementary Table S7.

### MSPDBL2 is expressed in schizonts independently of AP2-G expression

Given that GDV1 induces sexual commitment through de-repression of the epigenetically silenced *ap2-g* locus (22, 28), it was tested whether AP2-G is expressed in the same individual schizonts as MSPDBL2. Using an engineered parasite line with HA-tagged AP2-G (3D7/AP2G-HA) (29), schizonts were examined by two-colour immunofluorescence, using anti-HA antibody to detect AP2-G alongside antibodies to a conserved region of MSPDBL2. It was expected that proportions of schizonts positive for MSPDBL2 would be very low, as data presented above show < 1% of 3D7 schizonts express the protein. Out of a total of 5571 mature schizonts examined (each having at least 8 nuclei), only two schizonts were positive for both proteins, whereas 30 schizonts were positive for MSPDBL2 only (these were negative for AP2-G), and seven were positive for AP2-G only (these were negative for MSPDBL2) (Figure 5). This finding at the protein level is consistent with results from a single cell transcriptomic analysis of parasite line HB3, which indicated *ap2-g* and *mspdbl2* transcripts occurred mostly in different schizonts (17). The results indicate that, although AP2-G and MSPDBL2 expression are both upregulated by GDV1, the subsequent pathways of differentiation are independent so that MSPDBL2 is not a marker of sexually committed schizonts.

**Figure 5.**
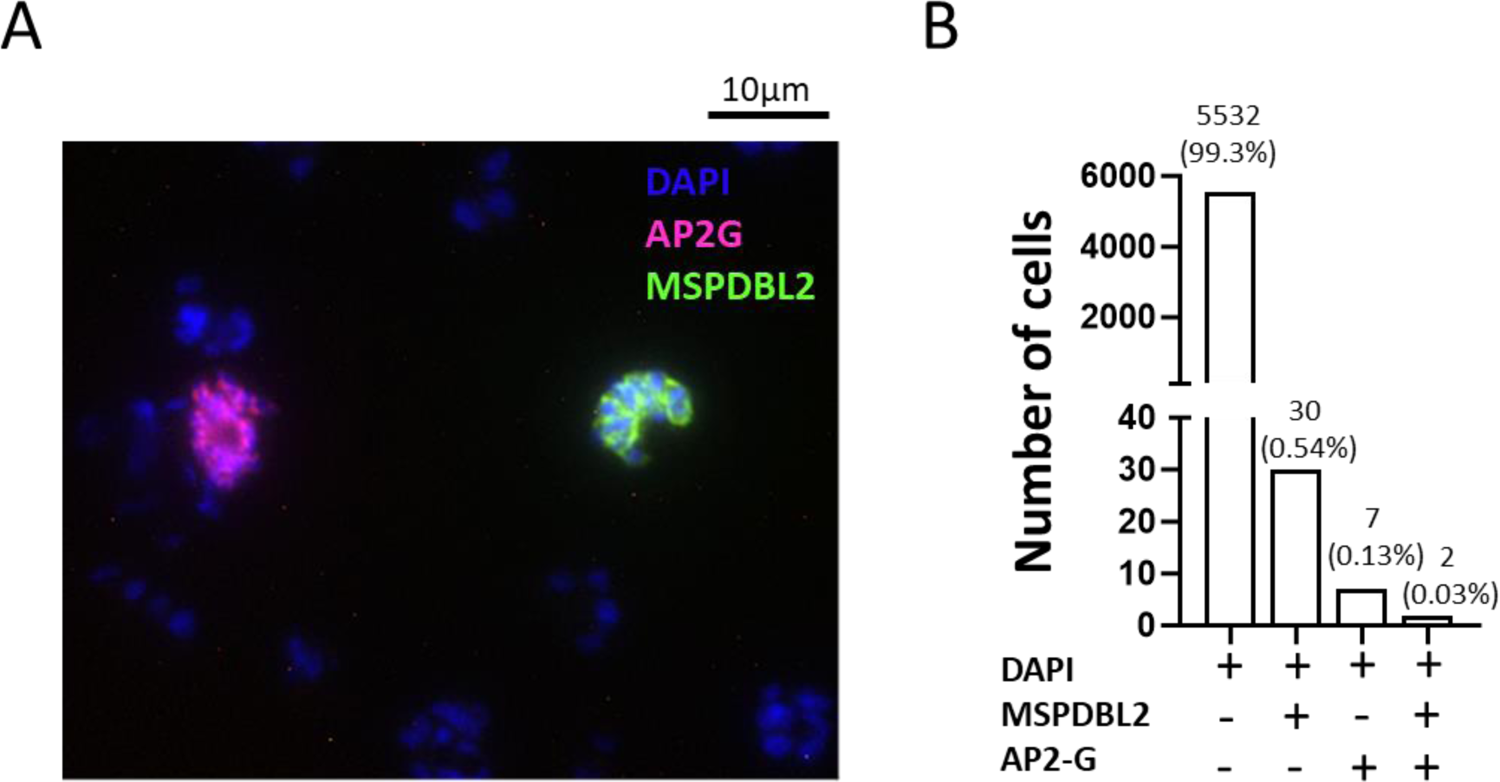
MSPDBL2 expression in individual schizonts is not associated with AP2-G expression. **A.** Immunofluorescence microscopy was performed on 3D7/AP2G-HA parasites which have an HA tag engineered on the endogenous AP2G. Staining was performed using anti-N-terminal MSPDBL2 murine Ab to detect MSPDBL2 (green), α-HA rat Ab to detect AP2-G (red), nuclei stained blue with DAPI). This microscopy image shows two schizonts that were positive, each for the separate respective proteins. **B**. Proportions of mature schizonts in the 3D7/AP2G-HA line expressing either MSPDBL2 alone, AP2G alone, or both markers. A total of 5571 mature schizonts with at least 8 nuclei were examined. Most schizonts were negative for both proteins (5532), but seven were positive for AP2G alone, 30 were positive for MSPDBL2 alone and two were positive for both markers.

### Generation of *Plasmodium falciparum* lines expressing a tagged or disrupted version of MSPDBL2 in a majority of schizonts

A strategy based on the selection-linked integration (SLI) system (30) was employed to generate parasite lines expressing a C-terminal TdTOM-3xHA-tagged full-length version of MSPDBL2 or 3xHA-tagged truncated version of MSPDBL2. The 3D7/iGP parasite line was chosen as the background for engineering, as it allows conditional overexpression of GDV1 and induction of elevated gametocyte conversion rates (23). The SLI system was used to enable modification at the endogenous *mspdbl2* locus to allow selection of schizonts expressing either an intact full-length MSPDBL2 (referred to as MSPDBL2-TAG) or a disrupted version of MSPDBL2 (referred as MSPDBL2 DEL) lacking two-thirds of the protein that covers the DBL domain (Figure 6A)(Supplementary Figures S1 and S2). It is important to note that no Shield-1 reagent was added at the selection stage, so there was no manipulation of GDV1 expression in that part of the process, but the engineered MSPDBL2-expressing parasites were selected for gentamycin-resistance.

**Figure 6.**
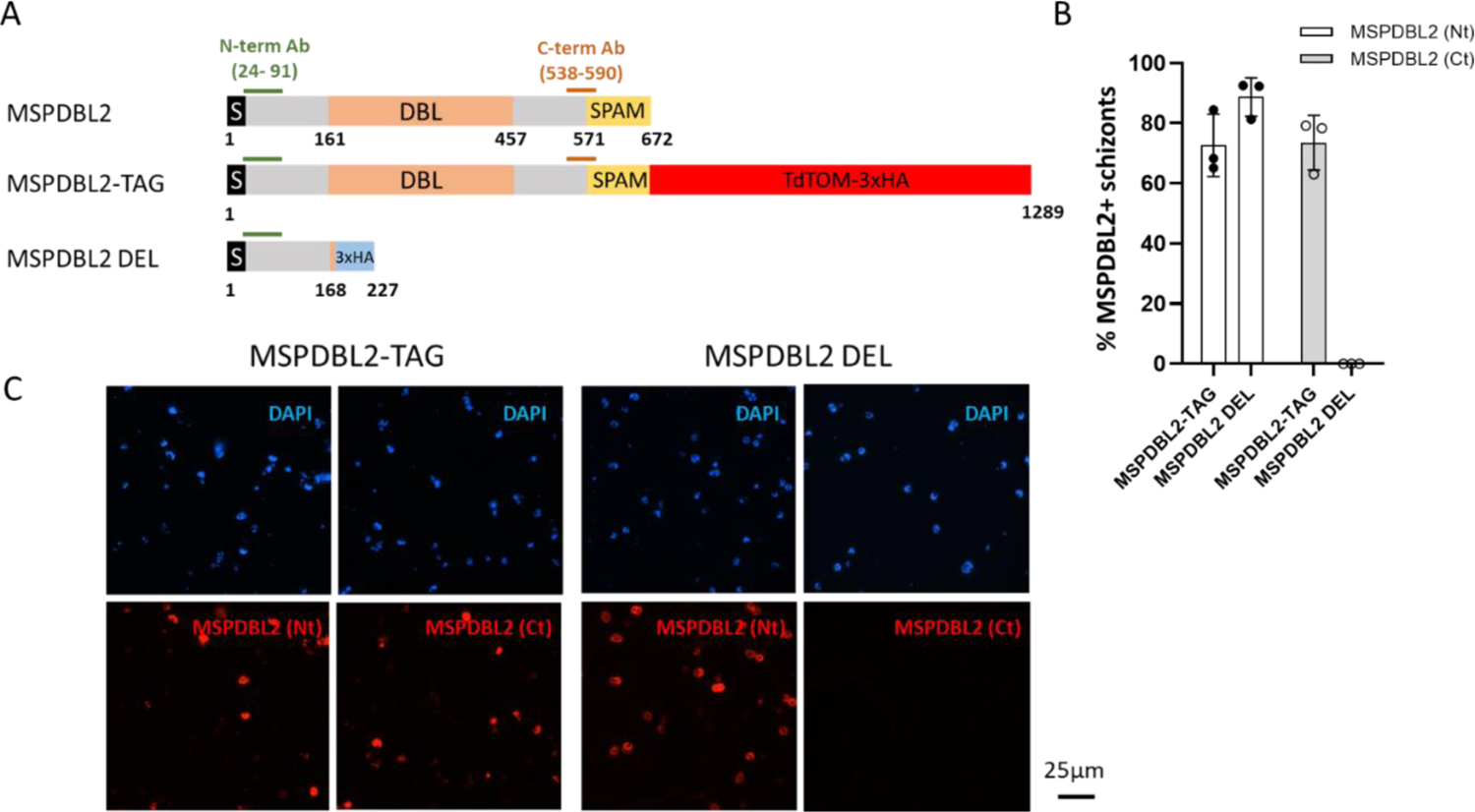
Generation of *P. falciparum* lines expressing intact or disrupted MSPDBL2 in a majority of schizonts. **A.** Schematic representation of the primary structural characteristics of MSPDBL2, as well as the TdTOM-3xHA tagged replacement (MSPDBL2-TAG) and the 3xHA-tagged disrupted version (MSPDBL2 DEL), both at the endogenous locus in the different selection linked integration (SLI) lines. The N-terminal signal sequence, the DBL (‘Duffy binding-like’) and SPAM (‘secreted polymorphic antigen associated with merozoites’) domains and the location of the recombinant protein sequences used to raise the α-N-terminal and α-C-terminal antibodies are indicated. **B.** Following G418 selection for gentamycin-resistant integrant parasites, most schizonts expressed either the entire or truncated version of MSPDBL2 in the respective lines, as assessed by immunofluorescence using antibodies against N-terminal or C-terminal MSPDBL2 sequences. All individual counts and assay values are given in Supplementary Table S8. **C.** Representative immunofluorescence (red) microscopy of schizonts expressing MSPDBL2 as an intact tagged version (MSPDBL2-TAG) or a disrupted version (MSPDBL2 DEL) stained by antibodies in murine sera against N-terminal (Nt) or C-terminal MSPDBL2 peptides (Ct). Parasite nuclei are stained with DAPI (blue).

Polyclonal antibodies against the conserved N-terminal region of MSPDBL2 (5) were used to detect either the intact or truncated protein, while polyclonal antibodies against the conserved C-terminal region of MSPDBL2 (5) were used to detect the intact protein (Figure 6A). After repeated rounds of G418 selection for parasites with the modified locus, a majority of schizonts in the MSPDBL2-TAG line expressed intact MSPDBL2 (staining with either the anti-N-terminal MSPDBL2 Ab or the anti-C-terminal MSPDBL2 Ab (Figure 6B and 6C, Supplementary Table S8). In comparison, the MSPDBL2 DEL line expressed a truncated version of MSPDBL2 in most schizonts (positive with the anti-N-terminal but not with the anti-C-terminal MSPDBL2 Ab) (Figure 6B and 6C, Supplementary Table S8).

### Expression of an intact or disrupted MSPDBL2 does not affect rates of sexual commitment

To test whether there was any direct effect of expressing intact or disrupted MSPDBL2 on the rates of gametocyte conversion per cycle, assays involving over-expression of GDV1 were conducted with the *mspdbl2*-engineered lines as described above. Each line was tested with multiple experimental replicates, in the absence and presence of Shield-1 reagent to stabilise the overexpression of GDV1, which drives sexual commitment.

Gametocyte conversion rates were assayed using a method previously described to detect Pfs16-positive parasites early in the following cycle (24). All parasite lines showed massively increased gametocyte conversion rates in response to overexpression of GDV1 under Shield-1 (Figure 7). There were no significant differences in the rates between MSPDBL2-TAG and MSPDBL2 DEL lines in which most parasites respectively express intact or disrupted MSPDBL2, and both showed similar rates to the 3D7/iGP parental line (Figure 7, Supplementary Table S9). This confirms that intact MSPDBL2 is not required for sexual conversion or subsequent sexual differentiation.

**Figure 7.**
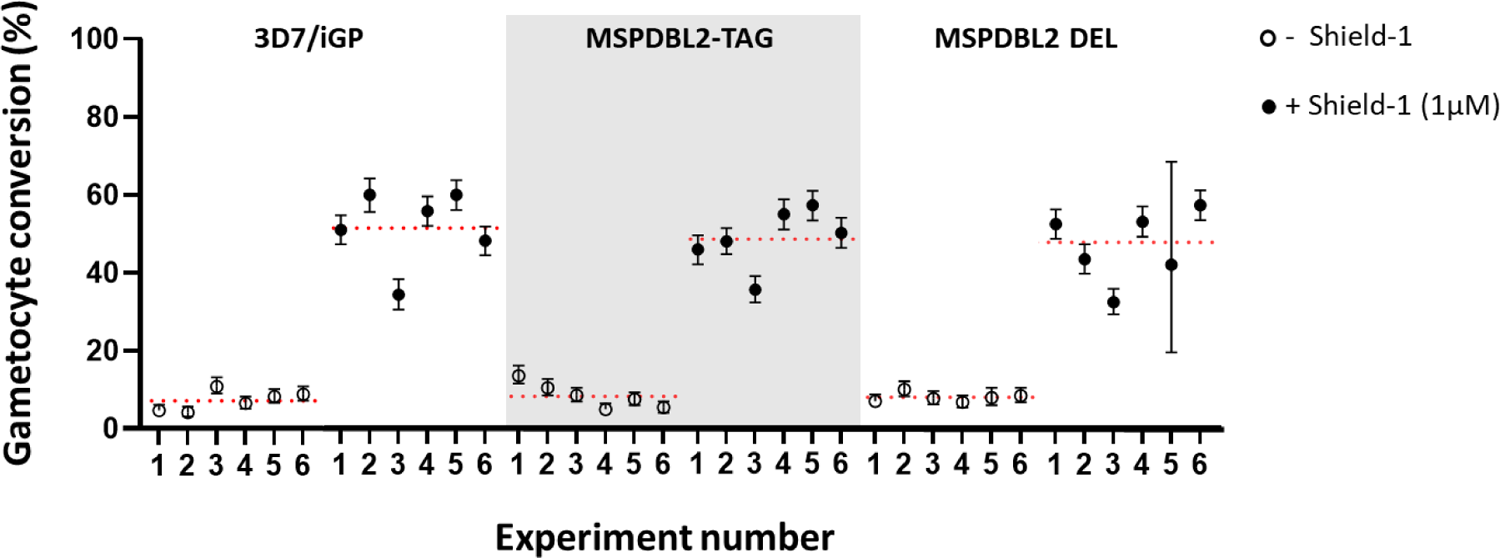
Sexual conversion and early gametocyte development is not affected by disruption or tagging of MSPDBL2. **A.** Gametocyte conversion rates of MSPDBL2-TAG and MSPDBL2 DEL lines compared to the parental 3D7/iGP line. Six experimental replicates were performed on each line in the presence and absence of 1 µM Shield-1 reagent to stabilize the overexpression of GDV1 (using the same protocol as described for assays in Figure 1). Red dotted lines represent the means across six replicate assays. All individual counts and assay values for all replicate assays are given in Supplementary Table S9.

### Expression of an intact or disrupted MSPDBL2 does not affect asexual multiplication rates

To test whether there was an effect of expressing intact or disrupted MSPDBL2 on asexual growth, the asexual blood stage multiplication rates of parasites were analysed, comparing the engineered lines in which most schizonts expressed the full length or disrupted version of MSPDBL2, as well as parasites in which most schizonts are MSPDBL2-negative. Each of the parasite lines was tested in three separate six-day exponential growth assays, each separate assay involving triplicate flasks with erythrocytes from different donors using a protocol described previously (31, 32). This showed that the multiplication rates of 3D7/iGP, MSPDBL2-TAG and MSPDBL2 DEL were not significantly different (Figure 8), all being approximately 8-fold per 48 hours. These rates were similar to those of the 3D7 parasite line that was incorporated alongside as a control for assay quality (Supplementary Table S10), which was consistent with its previously established multiplication rate using this assay method (31, 32).

**Figure 8.**
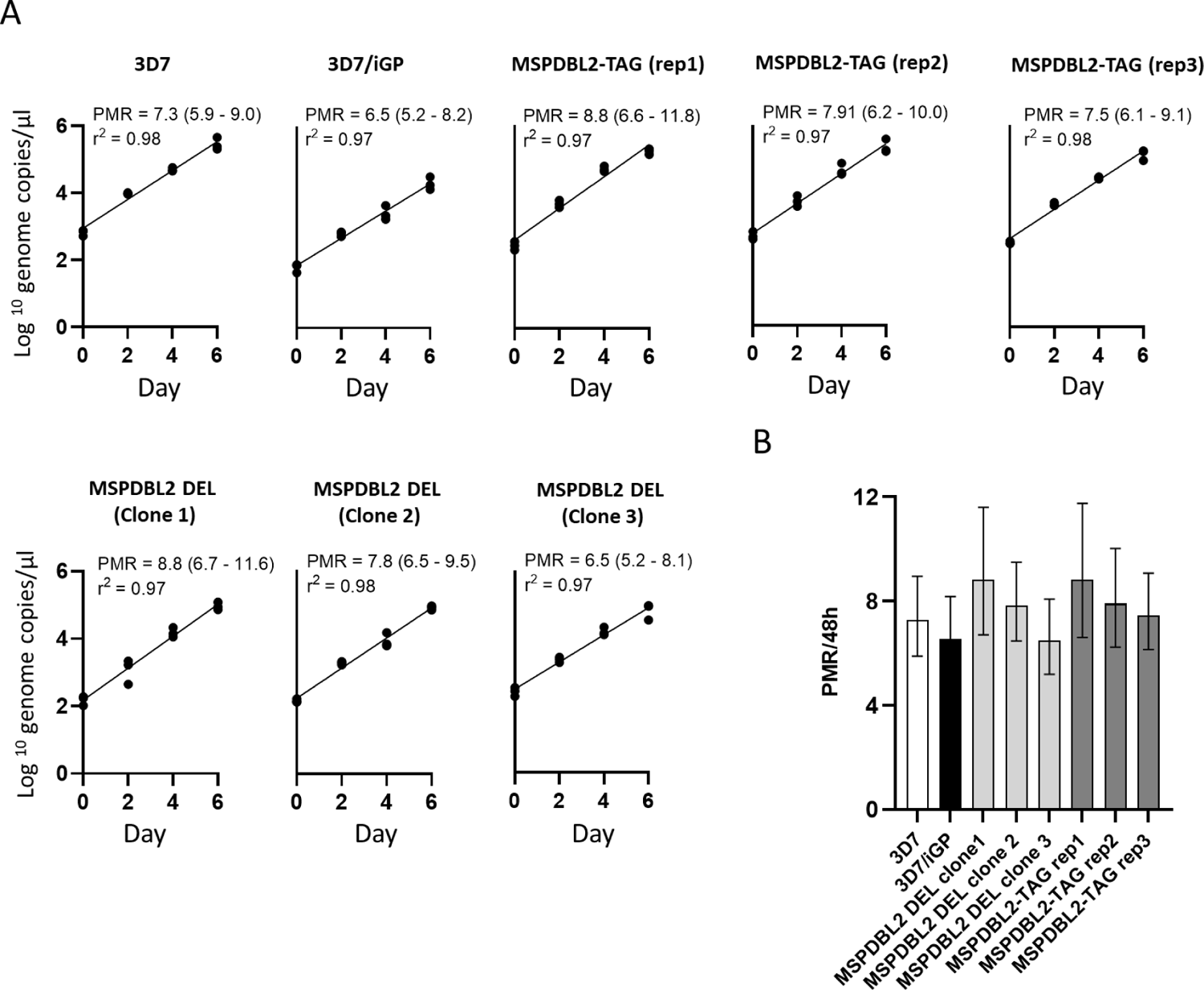
Asexual multiplication rates are not affected by expression of intact or disrupted MSPDBL2. Exponential multiplication rate assays performed over 6 days. Parasite density (log_10_ scale) is expressed as numbers of genome copies per μl of a standard volume of extracted DNA at each sampled timepoint, using quantitative PCR. Experiments were performed on three biological replicates of MSPDBL2-TAG, and three MSPDBL2 DEL clones, each compared to the parental 3D7/iGP line, and 3D7 which is always incorporated as a control (with previously known multiplication rate of approximately 8-fold per 48 hours). Each assay was performed in triplicate, using erythrocytes from different blood donors in separate flasks. **B**. Histograms presenting general linear model estimations (with 95% confidence intervals) of the multiplication rates per 48 hours over the duration of the assays. All numerical data are shown in Supplementary Table S10).

## DISCUSSION

This study shows that the *P. falciparum* regulator GDV1 is involved in more than one parasite differentiation process, not restricted to regulating sexual commitment, but also regulating proportions of schizonts expressing MSPDBL2 although most are asexually-committed. Using an engineered parasite line with overexpression of GDV1, there was a significant correlation between the MSPDBL2-positive schizont proportions and rates of gametocyte conversion, indicating that both AP2-G and MSPDBL2 are positively regulated by GDV1. This follows a previous observation that GDV1 over-expression led to increased *mspdbl2* transcript levels (22), in a study indicating that GDV1 is an upstream activator of sexual conversion through eviction of the repressor HP1 from the normally silenced *ap2-g* locus (22, 28). Along with *ap2-g*, *mspdbl2* is one of the few genes outside of sub-telomeric chromosomal regions to possess an H3K9me3 heterochromatic mark of gene silencing (14) targeted by HP1 (33),(15). It was previously considered that expression of MSPDBL2 may be a feature of schizonts committed to sexual development (16, 22), but the present study does not support this.

Surveying multiple independent culture preparations of a diverse panel of *P. falciparum* cultured lines showed varying proportions of MSPDBL2 positive schizonts, confirming and extending previous results on some of the lines (5). Across all the diverse parasite lines, or among replicates of individual lines, there was no significant correlation between the proportion of MSPDBL2 positive schizonts and commitment to develop into gametocytes in the following cycle. Interestingly, MSPDBL2 positive schizonts were detected in parasite line F12, a 3D7-derived clone containing a loss-of-function mutation in *ap2-g* (34, 35), which never produces gametocytes (24, 35). This extends a previous observation that GDV1 overexpression in the F12 background led to an increase in *mspdbl2* transcript expression (22), although these parasites do not undergo sexual development. Furthermore, the finding here that MSPDBL2 and AP2-G are mainly expressed in different individual schizonts, is consistent with a single-cell RNAseq analysis indicating that *mspdbl2* and *ap2-g* gene transcripts are mostly detected in different schizonts (17).

Previous analysis of mature *P. falciparum* schizonts during the first *ex vivo* cycle of development in clinical isolates showed a skewed distribution of MSPDBL2 positive schizont proportions similar to that seen among the long-term cultured parasite lines, with most isolates having < 1% schizonts positive but a minority having significantly higher proportions (17). In a bulk transcriptome analysis of a subset of those isolates, varying proportions of schizonts expressing MSPDBL2 correlated with relative transcript levels of many gametocyte-associated genes, although they did not correlate with *ap2-g* transcript levels. Correlation with *gdv1* transcript levels was not seen and would not have been expected in that study, as *gdv1* is regulated by antisense RNA competing with the coding sense strand (22, 36), whereas the transcriptome analysis was based on combined RNA-seq of either strand so it could not discriminate between positive and suppressive transcripts. In the present study, addition of choline to serum-free medium had a significant suppressive effect on proportions of schizonts expressing MSPDBL2 in the HB3 parasite line, consistent with the known effect of choline in suppressing *gdv1* expression (22). The physiological mechanisms whereby choline concentration affects parasite sexual commitment is an area of recent investigation (37, 38), which may also be of relevance for studying regulation of MSPDBL2.

As both AP2-G and MSPDBL2 are under the control of GDV1, it remains unknown how their expression is dissociated so that they appear in different mature schizonts. Given that the *mspdbl2* locus has a marked heterchromatin signature (14), once the expression of MSPDBL2 is achieved within a given cycle, active expression in schizonts in subsequent cycles may be epigenetically inherited even in the absence of renewed GDV1 expression. It has been shown that, once established, heterochromatin distribution and the active or silenced state of other clonally-variant genes are stably inherited through multiple asexual cycles (39). Parasites that activate AP2-G expression abandon asexual multiplication (40), so the effect of GDV1 on AP2-G is only within the same cycle, whereas MSPDBL2 expression may reflect GDV1-dependent activation events occurring in previous cycles. It will be relevant to discover either within-cycle or across-cycle mechanisms that explain how expression of MSPDBL2 is dissociated from AP2-G among different individual parasites, while recognizing that some of the regulatory processes may be probabilistic rather than entirely deterministic (1). Investigation of co-expression with other proteins may be relevant, as well as effects on proteins expressed very early in the following cycle such as GEXP05 which is upregulated by GDV1 (41) and AP2-G (29, 42) although it can also be expressed in AP2-G-null parasites (43).

Results also showed that selection for expression of either intact or truncated MSPDBL2 in a majority of schizonts, using an engineered selection-linked integration system, did not affect gametocyte conversion rates in culture, as the latter remained sensitive to overexpression of GDV1. It should be noted that these experiments involve an artificial system to engineer the allelic replacements under antibiotic selection at the endogenous *mspdbl2* locus, which might alter a potentially fragile heterochromatin domain at the locus. Future studies of chromatin accessibility and histone modification will be relevant for understanding regulation of processes that must occur after GDV1 expression, as these must determine which individual parasites will go on to express the MSPDBL2 antigen and which will become sexually committed. Beyond this, interpreting the function of the *mspdbl2* gene in natural infections requires further investigation. Despite strong inference indicating MSPDBL2-positive parasites are targets of protective immune responses (3, 5, 7, 44), so that there would be background selection against expression, the biological significance of this distinct antigenic subpopulation remains to be discovered.

It is plausible that MSPDBL2 may broaden or change the invasion pathway of merozoites, although its erythrocyte receptor is unknown (7, 12). Here, there was no significant difference in the overall exponential multiplication rate of parasite cultures in which most schizonts expressed either an intact or truncated MSPDBL2, compared to cultures in which most schizonts were negative. However, it is possible that MSPDBL2-positive parasites have a replication advantage in particular environments within human infections or might survive stress under some conditions experienced during infections. A hypothesis that these parasites have an enhanced ability to tolerate some unusual stress conditions may be supported by a previous observation that overexpression of MSPDBL2 can enable parasites to survive concentrations of some antimalarial drugs that are otherwise inhibitory (21). Under this hypothesis, stress conditions *in vivo* may induce upregulation of the MSPDBL2-positive subpopulation as a diversification to increase the likelihood that some asexual parasites survive, while another subpopulation of mostly MSPDBL2-negative parasites commit to gametocytogenesis and transmission.

## MATERIALS AND METHODS

### Plasmodium falciparum cultured lines

The inducible gametocyte producer *P. falciparum* clone 3D7/iGP_D9 was studied (23) (referred to as 3D7/iGP for brevity in the current paper), to assess the impact of GDV1 overexpression on the proportion of MSPDBL2+ schizonts using methods described in a separate subsection below. To test the effect of *gdv1* gene disruption on expression of MSPDBL2, the *P. falciparum line* NF54 GDV1 delta 39 was studied (this has a truncated *gdv1* coding sequence previously shown to be non-functional for promoting sexual commitment) (25) and compared with its parental line NF54 that has an intact *gdv1* gene. To analyse variation in proportions of MSPDBL2+ schizonts among isolates, and test for correlations with gametocyte conversion rates, thirteen other *P. falciparum* lines were also studied, with putative origins given here in parentheses: NF54 (an ‘airport malaria’ isolate from Netherlands with genetic similarities to African isolates), 3D7 (cloned from NF54), F12 (derived from 3D7 with a nonfunctional mutated *ap2-g*), D10 (Papua New Guinea), D6 (Sierra Leone), Dd2 (Southeast Asia), FCC2 (China), HB3 (Honduras), R033 (Africa), Palo Alto (Uganda), 7G8 (Brazil), GB4 (Ghana). All lines were thawed from pre-existing frozen stocks of cultures at the London School of Hygiene and Tropical Medicine and were not re-cloned prior to this study. To examine whether MSPDBL2 was expressed in the same schizonts as AP2-G, the parasite line 3D7/AP2G-HA (with a 3xHA epitope-tagged AP2-G engineered into the E5 clone of 3D7) (29) was used. Additional parasite modifications conducted in the present study are described in a separate sub-section below.

### Immunofluorescence assays to quantify proportions of mature schizonts expressing MSPDBL2

The *P. falciparum* lines were cultured as described previously (24), being maintained in RPMI 1640 media supplemented with AlbuMAX™ II in human erythrocytes at 3% hematocrit and incubated at 37°C in atmospheric air with 5% CO_2_. Late-stage parasites containing hemozoin were positively selected using MACS® LD magnetic columns (45).

Within 24 hours, this was followed by a second MACS purification, from which the flow through containing ring-stage parasites was collected, and returned into culture, following which parasites were allowed to develop until a large proportion were at schizont stages (24). Cells from these cultures were harvested by centrifugation and washed in PBS/3% BSA, resuspended to 2% hematocrit and spotted onto multiwell slides (Hendley, Essex, UK) which were then air-dried and stored at −80°C. Mature schizonts (with at least 8 nuclei) were analyzed by immunofluorescence microscopy using murine polyclonal serum specific for an N-terminal conserved region of MSPDBL2 (5, 17) to determine the proportions that were MSPDBL2+, as this gives clear positive and negative discrimination following a protocol previously described (5, 17).

Briefly, slides were fixed for 30 min with PBS/4% paraformaldehyde and permeabilized for 10 min with PBS/0.1% Triton X-100. The polyclonal anti N-terminal MSPDBL2 serum was diluted 1:200 in PBS/3% BSA and incubated for 30 minutes at room temperature. Subsequently, Alexa Fluor 594-conjugated anti-mouse IgG (Invitrogen, A11032) was used as secondary antibody, diluted 1:1000 in PBS/3% BSA and incubated for 30 minutes at room temperature. Finally, DAPI-containing ProLong^TM^ Diamond antifade mountant (ThermoFisher Scientific) was used as slides mounting media. Parasite counting was performed using a Zeiss CCD fluorescence microscope, identical settings were used for each experiment. Approximately 1000 parasites were counted and used to determine the proportions of mature schizonts expressing MSPDBL2 in each preparation, although occasionally lower numbers were counted if parasitemia was low.

For the parasite line 3D7/AP2G-HA in which the AP2-G is 3xHA epitope-tagged (29), two-colour immunofluorescence was performed to discriminate schizonts expressing MSPDBL2 and AP2-G. In this case, the polyclonal murine anti N-terminal MSPDBL2 serum and a rat monoclonal α-HA antibody (Roche 3F10) were used, with fluorescine-conjugated anti-mouse IgG and Alexa Fluor 594-conjugated anti-rabbit IgG secondary antibodies used along with DAPI staining.

To compare the proportion of mature schizonts expressing MSPDBL2 in medium with and without added choline, multiple replicate cultures of HB3 and NF54 parasites were prepared similarly as described above and then split after the second MACS column purification (used to isolate the ring stages contained in the flow through) into separate wells with a minimal culture medium (RPMI 1640 with 25 mM HEPES, 100 mM Hypoxanthine, 1 mM L-glutamine, 0.39% fatty acid-free BSA, 30 mM Palmitic acid, 30 mM Oleic acid) either supplemented with 2 mM choline or lacking choline. Parasites were exposed to this treatment from the ring stage of development onwards and allowed to grow until the schizont stage to be analysed by IFA as described above.

### MSPDBL2+ schizont proportions and sexual conversion rates in engineered parasite lines

The *P. falciparum* 3D7/iGP line was used to test the effect of GDV1 overexpression on the proportion of MSPDBL2+ schizonts. When parasites were at the ring stage, specified concentrations of Shield-1 reagent (from zero to 1 µM, Supplementary Tables S1 and S2) were added to culture media, as Shield-1 stabilizes the overexpressed GDV1-GFP-DD (the destabilization domain DD tagged to a GFP motif having been fused to GDV1 and integrated into the dispensable *cg6* locus *PF3D7_0709200*) (22). Parasites were allowed to grow until most were at trophozoite stages (30-35 hours post invasion), and the GDV1+ proportion was analyzed using an EVOS thermoFisher fluorescence microscope. A portion of the culture was harvested and stained with DAPI, and the GDV1+ proportion determined by differential counting of GFP positive trophozoites as a proportion of all DAPI positive trophozoites.

Then, the parasites were allowed to grow until the schizont stage, when a portion of the culture was harvested and analyzed to determine proportions of MSPDBL2+ schizonts among mature schizonts with at least 8 nuclei, as described above. The remaining parasite culture was allowed to re-invade erythrocytes, and after 38-46 hours parasites were analyzed for gametocyte conversion using Pfs16/DAPI staining as described previously (24). Briefly, a monoclonal α-Pfs16 murine antibody 93A3A2 (46) was diluted 1:2000 in PBS/3% BSA and incubated for 30 minutes at room temperature. Alexa Fluor 594-conjugated anti-mouse IgG (Invitrogen, A11032) was used as secondary antibody, diluted 1:1000 in PBS/3% BSA and incubated for 30 minutes at room temperature, and DAPI-containing ProLong^TM^ Diamond antifade mountant (ThermoFisher Scientific) was used. Parasite counting was performed using a Zeiss CCD fluorescence microscope, identical settings were used for each experiment. A minimum of 300 parasites were counted to determine the proportion of Pfs16 positive stage-I gametocytes.

### Generation of transgenic parasites expressing intact tagged or disrupted MSPDBL2

Tagging of MSPDBL2 with C-terminal TdTOM-3xHA for expression in a majority of schizonts (MSPDBL2-TAG line) was performed by modifying the endogenous *mspdbl2* locus through single homologous recombination using the selection-linked integration (SLI) T2A/G418 strategy (30). To construct the transfection vector, a 1002 bp region of *mspdbl2* starting 1284 bp downstream from the start codon and lacking the stop codon was amplified (primers P3 and P4, Supplementary Table S11) and inserted into a Xho1/BamH1 digested pL7M2_TdTom_T2A/G418 plasmid vector. This vector corresponds to a modified version of a pL7M2_T2A/G418 vector (47) with *TdTomato* sequence amplified from a pRHOPH3-RAP1-tdTom plasmid (48) inserted in phase of the N-terminal of the *3ha_t2a_neoR* (neomycin resistance) locus of the pL7M2_T2A/G418 vector previously digested with BamH1 and Kpn1 (primers P1 and P2, Supplementary Table S11, Supplementary Figure S1). To generate *mspdbl2*-disrupted parasites expressing a short truncated version of the protein (MSPDBL2 DEL line), the selection-linked integration (SLI) T2A/G418 strategy was employed, using a transfection plasmid with a truncated *mspdbl2* sequence (up to 505 bp from the start codon, amplified with primers P8 and P9, Supplementary Table S11) inserted in frame with a 3xHA tag and the N-terminal of the *3ha_t2a_neoR* (neomycin resistance) locus of the pL7M2_T2A/G418 vector previously digested with BamH1 and Kpn1 (47) (Supplementary Figure S2). The inserted modified *mspdbl2* sequences in the plasmid vector constructs are shown (Supplementary Table S12).

Both constructs were transfected into purified schizonts of the 3D7/iGP *P. falciparum* line (23) using an AMAXA nucleofector 4D (Lonza) and P3 reagent and selected as previously described with some modifications (30). Briefly, parasites were transfected with 40 µg of purified plasmid. 5nM of WR99210 (WR) was used to positively select the transfectants. Then, the transfected parasites were subjected to selection using 1 mg/ml G418 (Cambridge bioscience, 1557-1G) for 4-6 days to select integrants able to produce the tagged or truncated version of MSPDBL2. The cultures were then routinely subjected to additional rounds of G418 selection, with one round of selection every 4 to 5 weeks. Integration was monitored by PCR using specific primers to detect the 5’integration of the TdTOM-3xHA tag (Supplementary Figure S1) and detect the truncation in the disrupted line (Supplementary Figure S2) within the endogenous gene locus (Supplementary Figures S1 and S2, Supplementary Table S13).

The expression of MSPDBL2 in the tagged/disrupted lines was confirmed by immunofluorescence microscopy. To distinguish the intact and truncated MSPDBL2 expression, murine sera raised against the conserved N-terminal or C-terminal sequences region of MSPDBL2 were used as alternative primary antibodies for comparisons (1:200 in PBS/3% BSA) and incubated for 1 hour at room temperature. Following washing of slides, Alexa Fluor 594-conjugated anti-mouse IgG (1:1000 in PBS/3% BSA (Invitrogen, A11032)) was used as secondary antibody and incubated for 30 minutes at room temperature. DAPI-containing ProLong^TM^ Diamond antifade mountant (ThermoFisher Scientific) was used as mounting medium and parasite imaging was performed on an Inverted Nikon Ti Eclipse fluorescence microscope.

### Parasite multiplication rate assays

Parasite multiplication rate assays under exponential growth conditions were performed using a previously described protocol (31, 32). Erythrocytes were obtained from three anonymous donors, stored at 4°C until use within two days, and each assay was performed in triplicate by culturing with erythrocytes from each donor in a separate flask. Asynchronous parasite cultures were diluted to 0.02 % parasitaemia and 3 % haematocrit in 5 ml culture volumes, and grown over 6 days with 300 µl culture being collected for DNA extraction and qPCR every 48 hours (days 0, 2, 4, and 6), culture media being replaced on days 2, 4 and 5. To assess the numbers of parasite genome copies at each assay timepoint, qPCR analysis was performed targeting the highly conserved single locus *Pfs25* gene (31). Quality control filtering was performed to exclude any point with less than 1000 estimated genome copy numbers, any point either lower or more than 20-fold higher than measured for the same culture at the previous time point 2 days earlier, and any point among the triplicates with more than 2-fold difference from the others. An overall parasite multiplication rate (per 48 hours) was calculated for each assay with 95% confidence intervals using a standard linear model (Graphpad PRISM). The *P. falciparum* clone 3D7, previously determined to have an exponential multiplication rate of approximately 8-fold per 48 hours (31), was assayed in parallel as a control in all assays.

### Statistical analysis

Testing for the rank correlation between the proportion of MSPDBL2+ schizonts and gametocyte conversion rate was performed using Spearman’s ρ (rho) coefficient. Comparisons of MSPDBL2 + schizont proportions between different lines were analysed for significance using the non-parametric Mann-Whitney test, being performed on the multiple replicate assay measurements made for each line. For each individual biological replicate data point, the 95% confidence intervals in these proportions were calculated from the exact numbers of cells counted. Statistical analyses were performed using Prism version 9. All count data and other numbers are tabulated in the Supplementary Information file.

