## Supplementary for "Expression of the MSPDBL2 antigen in a discrete subset of *Plasmodium falciparum* schizonts is regulated by GDV1 but not linked to sexual commitment"

Supplementary Figures and Tables (Figures S1-S2, Tables S1-S12)

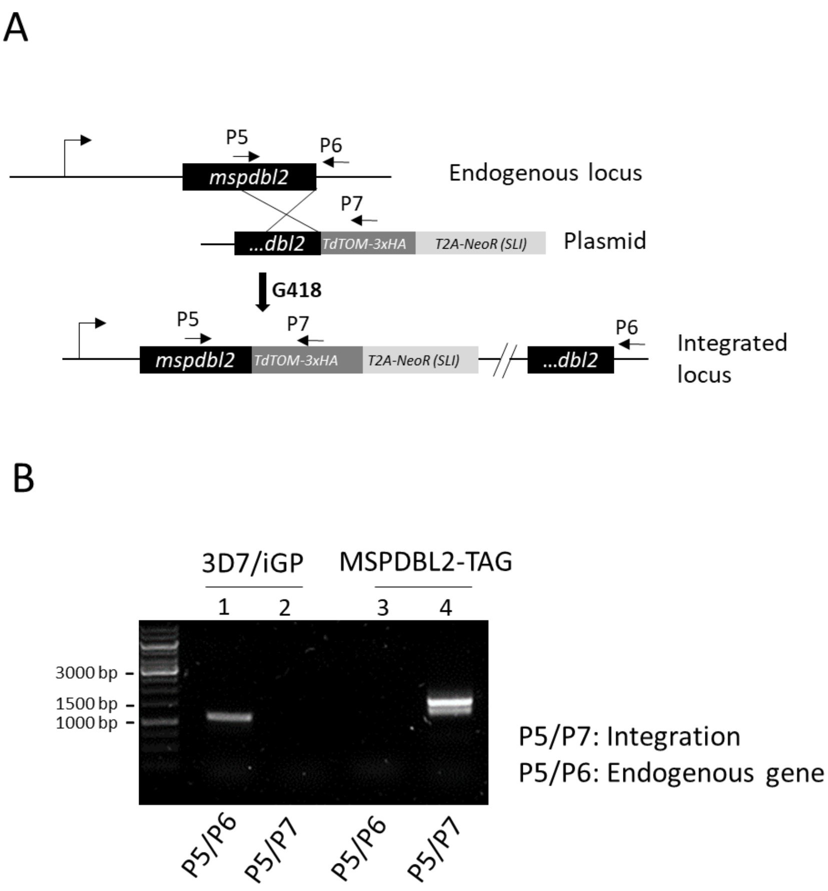


**Figure S1. Generation of a *Plasmodium falciparum* line to express a TdTOM-3xHA tagged version of MSPDBL2 in a majority of schizonts (MSPDBL2-TAG). (A)** Schematic representation of C-terminal tagging of MSPDBL2 with TdTOM-3xHA using the selection-linked integration (SLI-T2A/G418) single homologous recombination strategy. At the endogenous *mspdbl2* locus, the entire *mspdbl2* coding sequence (3D7 allele) is engineered in frame with TdTOM-3xHA and the *t2a* peptide linked to the *neomycin* selection marker (the contiguous engineered DNA sequence is shown in Table S13). Transgenic parasites were selected using G418. The position of the primers used for diagnostic PCR (listed in Table S12) are indicated by arrows. **(B)** Diagnostic PCR of MSPDBL2-TAG (lanes 3-4) compared to the parental 3D7/iGP line (lanes 1-2). Lanes 1 and 3: endogenous (P5-P6). Lanes 2 and 4: 5'integration (P5 and P7).

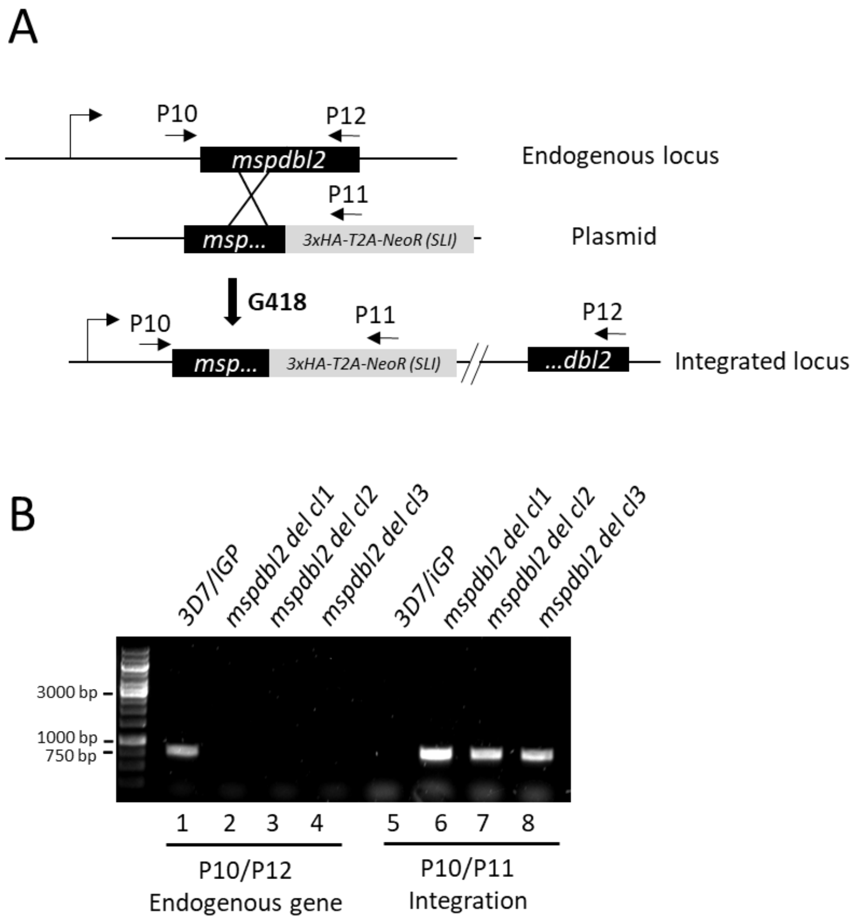


**Figure S2. Generation of a *Plasmodium falciparum* line to express a disrupted version of MSPDBL2 in a majority of schizonts (MSPDBL2 DEL lines). (A)** Schematic representation of the *mspdbl2* disruption strategy using selection-linked integration (SLI-T2A/G418) single homologous recombination. At the endogenous *mspdbl2* locus, the truncated version of *mspdbl2* is engineered in frame with and expressed as a 3xHA-tagged fusion containing the *t2a* peptide linked to the *neomycin* selection marker (the contiguous engineered DNA sequence is shown in Table S13). Transgenic parasites were selected using G418. The position of the primers (Table S12) used for diagnostic PCR are indicated by arrows. **(B)** Diagnostic PCR of MSPDBL2 DEL: Lanes 1-4: detection of the endogenous *mspdbl2* (P10-P12). Lanes 5-8: *mspdbl2* gene disruption (P10-P11). Lanes 1 and 5: 3D7/iGP. Lanes 2 and 6: MSPDBL2 DEL clone 1. Lanes 3 and 7: MSPDBL2 DEL clone 2. Lanes 4 and 8: MSPDBL2 DEL clone 3.

**Table S1.** Immunofluorescence microscopy counts of proportions of MSPDBL2 +ve schizonts among all those with at least 8 nuclei, for the 3D7/iGP parasite line in presence of 0 or 1 mM of Shield-1 reagent (GDV1 is overexpressed and stabilized in the presence of Shield-1). Also shown are the proportions of GDV1 +ve trophozoites. Four replicate experiments were performed.

| Assay Number | [Shield-1] (μM) | GDV1 +ve/ DAPI +ve | Proportion of GDV1+ trophozoites (± 95%CI) | MSPDBL2 +ve/ DAPI +ve | Proportion of MSPDBL2+ schizonts (± 95%CI) |
|---|---|---|---|---|---|
| 1 | 0 | 7/107 | **6.5** (3.0 – 13.1) | 4/694 | **0.6** (0.2 – 1.5) |
| | 1 | 85/102 | **83.3** (74.8 – 89.4)* | 289/612 | **47.2** (43.3 – 51.2)* |
| 2 | 0 | 4/310 | **1.3** (0.4 – 3.4) | 12/646 | **1.8** (1.0 – 3.3) |
| | 1 | 239/300 | **79.7** (74.7 – 83.8)* | 275/653 | **42.1** (38.4 – 45.9)* |
| 3 | 0 | 5/131 | **3.8** (1.4 - 8.8) | 9/1009 | **0.9** (0.4 – 1.7) |
| | 1 | 171/207 | **82.6** (76.8 – 87.2)* | 277/871 | **31.8** (28.8 – 34.9)* |
| 4 | 0 | 0/300 | **0.0** | 12/987 | **1.2** (0.7 – 2.1) |
| | 1 | 183/310 | **59.03** (53.3-64.3)* | 246/753 | **32.7** (29.4 – 36.1)* |
| **Mean** | **0** | | **2.9** | | **1.1** |
| | **1** | | **76.2** | | **38.5** |

*:Fisher exact test p value: p < 0.00001

**Table S2.** Immunofluorescence microscopy counts of proportions of MSPDBL2 +ve schizonts among all those with at least 8 nuclei, in the 3D7/iGP parasite line with varying concentrations of Shield-1 reagent (GDV1 is overexpressed and stabilized in the presence of Shield-1). Also shown are the gametocyte conversion rates (GCR) measured in the same cultures as the proportion of stage 1 gametocytes developing in the first cycle of re-invasion using anti-Pfs16 staining (GCR data from Stewart *et al*, *Microbiology Spectrum* 2022).

| **Assay number** | **Concentration Shield-1** | **MSPDBL2 +ve** | **DAPI +ve** | **Proportion of MSPDBL2+ schizonts (95%CI)** | **Pfs16 +ve** | **DAPI +ve** | **Pfs16 GCR (95%CI) *** |
|---|---|---|---|---|---|---|---|
| 1 | 0 | 7 | 868 | **0.8** (0.4 – 1.7) | 104 | 1171 | **8.9** (7.4 – 10.6) |
| | 0.055 | 310 | 1031 | **30.1** (27.3 – 32.9) | 157 | 1104 | **14.2** (12.3 – 16.4) |
| | 0.15 | 163 | 595 | **27.4** (24.0 – 31.1) | 188 | 1117 | **16.8** (14.7 – 19.1) |
| | 0.5 | 253 | 746 | **33.9** (30.6 – 37.4) | 197 | 1131 | **17.4** (15.3 – 19.7) |
| | 1 | 383 | 1046 | **36.6** (33.7 – 39.6) | 262 | 978 | **26.8** (24.11 – 29.65) |
| 2 | 0 | 7 | 877 | **0.8** (0.3 – 1.7) | 41 | 926 | **4.4** (3.3 – 6.0) |
| | 0.055 | 66 | 938 | **7.0** (5.5 – 8.9) | 164 | 800 | **20.5** (17.9 – 23.4) |
| | 0.15 | 127 | 882 | **14.4** (12.2 – 16.9) | 139 | 864 | **16.1** (13.8 – 16.7) |
| | 0.5 | 186 | 765 | **24.3** (21.4 – 27.5) | 209 | 790 | **26.5** (23.5 – 29.6) |
| | 1 | 159 | 840 | **18.9** (46.4 – 21.7) | 150 | 595 | **25.2** (21.9 – 28.8) |
| 3 | 0 | 12 | 987 | **1.2** (0.7 – 2.1) | 47 | 1043 | **4.5** (3.4 – 5.9) |
| | 0.055 | 77 | 922 | **8.3** (6.7 – 10.3) | 70 | 788 | **8.8** (7.1 – 11.1) |
| | 0.15 | 150 | 849 | **17.7** (15.2 – 20.4) | 140 | 629 | **22.3** (19.2 – 25.7) |
| | 0.5 | 193 | 806 | **23.9** (21.1 – 27.0) | 273 | 730 | **37.4** (34.0 – 41.0) |
| | 1 | 246 | 753 | **32.7** (29.4 – 36.1) | 267 | 701 | **38.1** (34.6 – 41.7) |

**Table S3.** Immunofluorescence microscopy counts of proportions of MSPDBL2 +ve schizonts among all those with at least 8 nuclei, in the NF54 GDV1Δ39 and NF54 parasite lines. The proportion of MSPDBL2+ schizonts are shown together with the 95% CI.

| **Assay number** | **Line** | **% Parasitemia** | **MSPDBL2 +ve** | **DAPI +ve** | **Proportion of MSPDBL2+ Schizonts (95%CI)** | **Fisher Exact test (p value)** |
|---|---|---|---|---|---|---|
| 1 | NF54 | 3.0 | 13 | 986 | 1.3 (0.7-2.3) | 0.0002 |
| | NF54 GDV1Δ39 | 1.8 | 0 | 1000 | 0 | |
| 2 | NF54 | 8.8 | 6 | 993 | 0.6 (0.24-1.35) | 0.031 |
| | NF54 GDV1Δ39 | 6.6 | 0 | 999 | 0 | |
| 3 | NF54 | 4.0 | 12 | 988 | 1.2 (0.67-2.14) | 0.0005 |
| | NF54 GDV1Δ39 | 6.5 | 0 | 1000 | 0 | |
| **Mean** | **NF54** | | | | **1.0** | |
| | **NF54 GDV1Δ39** | | | | **0** | |

**Table S4.** Immunofluorescence microscopy counts of proportions of MSPDBL2 +ve schizonts among all those with at least 8 nuclei, for 13 *P. falciparum* cultured lines with multiple culture replicates (74 preparations in total). The proportion of MSPDBL2+ schizonts are shown together with the 95% CI, and are matched with associated gametocyte conversion rate (GCR) data for 71 of the cultures (see footnote). For most of the replicates, the estimated parasitemia in the cycle during which the schizonts were tested for expression of MSPDBL2 is shown (% Para), as well as the next cycle parasitemia when the proportions of gametocytes were tested (% Para next cycle), and the fold change in parasitemia between these successive cycles (see footnote).

| Line | Replicate | RBC batch | % Para | MSPDBL2 +ve | DAPI +ve | Proportion of MSPDBL2+ schizonts (95% CI) | % Para next cycle | Fold change | Pfs16 GCR (95% CI) |
|---|---|---|---|---|---|---|---|---|---|
| F12 | 1 | 3 | - | 2 | 997 | 0.2 (0.0 - 0.8) | - | - | 0 |
| | 2 | 4 | - | 2 | 997 | 0.2 (0.0 – 0.8) | - | - | 0 |
| | 3 | 10 | 1.3 | 8 | 990 | 0.8 (0.4 – 1.6) | 8.7 | 6.7 | 0 |
| | 4 | 11 | 1.2 | 12 | 987 | 1.2 (0.7 – 2.1) | 5.1 | 4.3 | 0 |
| | 5 | 11 | 1.3 | 11 | 413 | 2.6 (1.4 – 4.8) | 2.2 | 1.7 | 0 |
| | ***Mean (SD)*** | | | | | ***0.9 (0.9)*** | | | ***0*** |
| D10 | 1 | 2 | - | 1 | 999 | 0.1 (0.0 – 0.6) | - | | 0.1 (0 - 5.6) |
| | 2 | 2 | - | 4 | 1994 | 0.2 (0.1 – 0.5) | - | | 0.8 (0.5 – 1.6) |
| | 3 | 3 | - | 1 | 999 | 0.1 (0.0 – 0.6) | - | | 0.2 (0.1 – 0.7) |
| | 4 | 9 | 3.4 | 0 | 1000 | 0.0 (0.0 - 0.5) | 8.8 | 2.6 | 0.3 (0.1 – 0.9) |
| | ***Mean (SD)*** | | | | | ***0.1 (0.1)*** | | | ***0.35 (0.31)*** |
| T9/96 | 1 | 13 | 1.1 | 0 | 999 | 0.0 (0.0 – 0.5) | 5.3 | 4.8 | 0.3 (0.1 – 1.0) |
| | 2 | 13 | 1.5 | 8 | 991 | 0.8 (0.4 - 1.6) | 4.5 | 3.0 | 0.3 (0.1 – 0.9) |
| | 3 | 13 | 2.2 | 3 | 996 | 0.3 (0.1 - 0.9) | 7.6 | 3.5 | 0.3 (0.1 – 1.0) |
| | 4 | 14 | 0.6 | 10 | 984 | 1.0 (0.5 - 1.9) | 12.5 | 21 | 0.2 (0.2 – 0.7) |
| | 5 | 14 | 0.6 | 2 | 997 | 0.2 (0.0 - 0.8) | 7.8 | 13 | 0.7 (0.2 – 2.0) |
| | ***Mean (SD)*** | | | | | ***0.5 (0.4)*** | | | ***0.36 (0.19)*** |
| Palo Alto | 1 | 6 | 0.8 | 2 | 303 | 0.6 (0.0 – 2.5) | 1.0 | 1.3 | 1.2 (0.7 – 2.1) |
| | 2 | 7 | 1.8 | 7 | 953 | 0.7 (0.3 – 1.5) | 2.7 | 1.5 | 2.7 (1.6 – 4.5) |
| | 3 | 7 | 1.7 | 1 | 998 | 0.1 (0.0 – 0.6) | 2.2 | 1.3 | 1.6 (0.7 – 3.6) |
| | 4 | 9 | - | 4 | 944 | 0.4 (0.1 – 1.1) | - | - | 0.1 (0 – 0.6) |
| | 5 | 9 | 1.4 | 0 | 383 | 0.0 (0.0 – 1.2) | 2.9 | 2.1 | 0.3 (0.1 – 0.9) |
| | 6 | 9 | 0.6 | 1 | 332 | 0.3 (0.0 – 1.9) | 1.4 | 2.3 | 0.1 (0 – 0.6) |
| | ***Mean (SD)*** | | | | | ***0.4 (0.3)*** | | | ***1 (1.03)*** |
| 3D7 | 1 | 2 | - | 0 | 329 | 0.0 (0.0 – 1.4) | - | - | 2.0 (1.3 – 3.1) |
| | 2 | 3 | - | 1 | 999 | 0.1 (0.0 – 0.6) | - | - | 0.3 (0.1 – 0.9) |
| | 3 | 3 | - | 2 | 997 | 0.2 (0.0 – 0.8) | - | - | 1.2 (0.7 – 2.1) |
| | 4 | 4 | 0.8 | 2 | 1996 | 0.1 (0.0 - 0.4) | 2.6 | 3.3 | 2.0 (1.3 – 3.1) |
| | 5 | 4 | - | 1 | 1080 | 0.1 (0.0 – 0.6) | - | - | 1.9 (1.2 – 3.0) |
| | 6 | 12 | 0.9 | 1 | 422 | 0.2 (0.0 – 1.5) | 1.1 | 1.2 | 2.3 (1.5 – 3.5) |
| | 7 | 12 | 1.1 | 8 | 860 | 0.9 (0.4 – 1.8) | 1.4 | 1.3 | 1.6 (1.0 – 2.6) |
| | ***Mean (SD)*** | | | | | ***0.2 (0.3)*** | | | ***1.6 (0.67)*** |

| Line | Replicate | RBC batch | % Para. | MSPDBL2 +ve | DAPI +ve | Proportion of MSPDBL2+ Schizonts (95% CI) | % Para. next cycle | Fold change | Pfs16 GCR* (95% CI) |
|---|---|---|---|---|---|---|---|---|---|
| | 1 | 5 | - | 14 | 543 | 2.6 (1.5 – 4.3) | - | - | 9.0 (7.3 - 11.1) |
| | 2 | 5 | - | 29 | 735 | 3.9 (2.7 – 5.6) | - | - | 1.6 (1.0 – 2,6) |
| **D6** | 3 | 5 | - | 26 | 974 | 2.6 (1.8 – 3.9) | - | - | 0.7 (0.3 – 1.5) |
| | 4 | 9 | 1.0 | 2 | 997 | 0.2 (0.0 – 0.8) | 0.1 | 0.1 | 2.7 (1.8 – 3.9) |
| | 5 | 9 | 1.4 | 7 | 647 | 1.1 (0.5 – 2.3) | 0.7 | 0.5 | 5.3 (4.0 – 6.9) |
| | ***Mean (SD)*** | | | | | ***2.1 (1.4)*** | | | ***3.8 (3.3)*** |
| | 1 | 6 | 1.4 | 28 | 972 | 2.8 (2.0 – 4.1) | 3.3 | 2.4 | 2.2 (1.5 – 3.4) |
| | 2 | 6 | 0.8 | 12 | 987 | 1.2 (0.7 – 2.1) | 1.5 | 1.9 | 9.3 (7.6 – 11.4) |
| | 3 | 6 | 2.5 | 36 | 729 | 4.9 (3.6 – 6.8) | 1.5 | 0.6 | 10.0 (8.3 – 12.2) |
| **GB4** | 4 | 6 | 1.2 | 13 | 866 | 1.5 (0.8 – 2.6) | 1.0 | 0.8 | 3.8 (2.8 – 5.3) |
| | 5 | 7 | 0.9 | 35 | 823 | 4.2 (3.1 – 5.9) | 7.7 | 8.6 | 6.3 (4.9 – 8.0) |
| | 6 | 7 | 1.4 | 133 | 866 | 15.4 (13.1 – 17.9) | 3.6 | 2.6 | 1.6 (1.0 – 2.6) |
| | 7 | 9 | 0.9 | 172 | 829 | 20.7 (18.1 – 23.6) | 2.2 | 3.7 | 3.5 (2.5 – 4.9) |
| | ***Mean (SD)*** | | | | | ***7.3 (7.1)*** | | | ***5.24 (3.36)*** |
| | 1 | 2 | 0.9 | 6 | 1995 | 0.3 (0.1 – 0.7) | 3.7 | 4.1 | 14.4 (12.3 -16.9) |
| | 2 | 3 | 1.5 | 1 | 95 | 1.0 (0.0 – 6.3) | 1.9 | 1.6 | 0.4 (0.2 – 1.00 |
| **Dd2** | 3 | 4 | - | 1 | 104 | 1.0 (0.0 – 5.7) | - | - | 4.8 (3.6 – 6.4) |
| | 4 | 4 | - | 4 | 995 | 0.4 (0.1 – 1.1) | - | - | 9.2 (7.5 – 11.2) |
| | 5 | 12 | - | 0 | 380 | 0.0 (0.0 – 1.2) | - | - | 4.3 (2.9 – 5.5) |
| | 6 | 12 | - | 21 | 980 | 2.1 (1.4 – 3.3) | - | - | 4.0 (2.9 – 5.5) |
| | ***Mean (SD)*** | | | | | ***0.8 (0.7)*** | | | ***6.1 (4.9)*** |
| | 1 | 2 | - | 69 | 467 | 14.8 (11.8 – 18.3) | - | - | 7.2 (5.5 – 9.4) |
| **HB3** | 2 | 2 | - | 107 | 892 | 12.0 (10.0 – 14.3) | - | - | 10.3 (8.6 – 12.3) |
| | 3 | 9 | 3.4 | 119 | 880 | 13.5 (11.4 – 16.0) | 4.5 | 1.3 | 6.1 (4.7 – 7.8) |
| | 4 | 9 | 0.9 | 67 | 535 | 12.5 (10.0 – 15.6) | 7.5 | 8.3 | 5.9 (4.6 – 7.6) |
| | ***Mean (SD)*** | | | | | ***13.2 (1.2)*** | | | ***7.3 (2.03)*** |
| | 1 | 2 | - | 17 | 983 | 1.7 (1.1 – 2.8) | - | - | 22.0 (19.6 – 24.7) |
| **NF54** | 2 | 2 | - | 7 | 992 | 0.7 (0.3 – 1.5) | - | - | 18.0 (15.7 –20.5) |
| | 3 | 13 | - | 0 | 150 | 0.0 (0.0 – 3.0) | - | - | 2.8 (1.9 – 4.0) |
| | ***Mean (SD)*** | | | | | ***0.8 (0.8)*** | | | ***14.2 (10.1)*** |

| Line | Replicate | RBC batch | % Para. | MSPDBL2 +ve | DAPI +ve | Proportion of MSPDBL2+ Schizonts (95% CI) | % Para. next cycle | Fold change | Pfs16 GCR * (95% CI) |
|---|---|---|---|---|---|---|---|---|---|
| | 1 | 5 | - | 2 | 340 | 0.6 (0.0 – 2.3) | - | - | 17.8 (15.4 – 20.4) |
| | 2 | 5 | - | 1 | 494 | 0.2 (0.0 – 1.3) | - | - | 7.2 (5.7 – 9.0) |
| | 3 | 7 | 1.3 | 10 | 989 | 1.0 (0.5 – 1.9) | 1.1 | 0.8 | 7.5 (6.0 – 9.4) |
| **RO33** | 4 | 7 | 2.4 | 0 | 422 | 0.0 (0.0 – 1.1) | 0.7 | 0.3 | 10.1 (8.3 – 12.2) |
| | 5 | 10 | - | 0 | 999 | 0.0 (0.0 – 0.5) | - | - | 7.8 (6.3 – 9.7) |
| | 6 | 10 | 1.4 | 2 | 349 | 0.6 (0.0 – 2.2) | 3.6 | 2.6 | 14.2 (12.0 – 16.6) |
| | ***Mean (SD)*** | | | | | ***0.4 (0.4)*** | | | ***10.7 (4.3)*** |
| | 1 | 6 | 2.1 | 0 | 1000 | 0.0 (0.0 – 0.5) | 3.2 | 2.7 | 18.6 (16.1 – 21.3) |
| | 2 | 6 | 1.2 | 0 | 474 | 0.0 (0.0 – 1.0) | 2.5 | 2.1 | 24.8 (22 – 27.9) |
| **7G8** | 3 | 10 | 1.9 | 3 | 332 | 0.9 (0.2 - 2.7) | 0.9 | 1.7 | 7.1 (4.8 – 10.3) |
| | 4 | 10 | 2.2 | 2 | 545 | 0.3 (0.0 – 1.4) | 2.6 | 1.2 | 2.0 (1.0 -3.7) |
| | 5 | 11 | 1.5 | 1 | 493 | 0.2 (0.0 – 1.3) | 2.2 | 1.5 | 4.8 (3.6 – 6.4) |
| | ***Mean (SD)*** | | | | | ***0.3 (0.3)*** | | | ***11.5 (9.7)*** |
| | 1 | 5 | 2.1 | 52 | 1257 | 4.1 (3.2 – 5.4) | 4.3 | 2.0 | 26.9 (23.9 – 30.1) |
| | 2 | - | - | 18 | 1184 | 1.5 (0.9 – 2.4) | - | - | 23.5 (20.0 – 26.6) |
| | 3 | - | - | 2 | 1336 | 0.1 (0.1 – 0.6) | - | - | 18.6 (16.1 – 21.4) |
| **FCC2** | 4 | 5 | 2.5 | 43 | 956 | 4.5 (3.3 – 6.0) | 3.9 | 1.6 | 17.6 (15.2 – 20.3) |
| | 5 | 5 | 1.1 | 4 | 934 | 0.4 (0.1 – 1.1) | 4.8 | 4.4 | 3.3 (2.0 – 5.3) |
| | 6 | 5 | 0.8 | 2 | 326 | 0.6 (0.0 – 2.4) | 3.0 | 3.8 | 4.6 (3.1 – 6.8) |
| | 7 | 10 | - | 3 | 996 | 0.3 (0.1 – 1.0) | - | - | 7.4 (5.9 – 9.2) |
| | 8 | 10 | - | 3 | 996 | 0.3 (0.1 – 0.9) | - | - | 4.9 (3.7 – 6.5) |
| | ***Mean (SD)*** | | | | | ***1.5 (1.8)*** | | | **13.3 (9.3)** |

The GCR data for these cultures were reported previously (Stewart et al. 2022 *Microbiology Spectrum* https://doi.org/10.1128/spectrum.02234-22) and are aligned here to enable analysis.

The parasitemia estimates were made for culture maintenance and are not available for all replicates, but those available are included here for *post hoc* exploratory analysis (this shows no correlation between these variables and MSPDBL2 expression in schizonts).

**Table S5.** Mann Whitney pairwise comparisons of proportions of MSPDBL2+ schizonts in different *P. falciparum* cultured lines. Statistically significant comparisons are highlighted in green (P<0.05*, P<0.01**, P<0.0001***), analysing data in Supplementary Table S3. Significant comparisons carried out with NF54 (n=3) were non-significant with every line tested except for GB4 (P=0.048).

| Lab Isolates | F12 | D10 | T9/96 | Palo Alto | 3D7 | D6 | GB4 | Dd2 | HB3 | RO33 | 7G8 |
|---|---|---|---|---|---|---|---|---|---|---|---|
| F12 | | | | | | | | | | | |
| D10 | 0.019* | | | | | | | | | | |
| T9/96 | 0.454 | 0.182 | | | | | | | | | |
| Palo Alto | 0.290 | 0.186 | 0.762 | | | | | | | | |
| 3D7 | 0.026* | 0.865 | 0.240 | 0.277 | | | | | | | |
| D6 | 0.238 | 0.024* | 0.063 | 0.052 | 0.008** | | | | | | |
| GB4 | 0.003** | 0.004** | 0.002** | 0.001*** | 0.001*** | 0.065 | | | | | |
| Dd2 | 0.961 | 0.081 | 0.398 | 0.372 | 0.053 | 0.125 | 0.003** | | | | |
| HB3 | 0.009** | 0.029* | 0.016* | 0.009** | 0.002** | 0.016* | 0.214 | 0.009** | | | |
| RO33 | 0.288 | 0.395 | 0.693 | 0.939 | 0.511 | 0.035* | 0.001*** | 0.387 | 0.009** | | |
| 7G8 | 0.167 | 0.587 | 0.587 | 0.654 | 0.866 | 0.04* | 0.002** | 0.173 | 0.016* | 0.688 | |
| FCC2 | 0.731 | 0.008* | 0.353 | 0.281 | 0.006** | 0.594 | 0.014* | 0.754 | 0.004** | 0.215 | 0.116 |

**Table S6.** Tests for correlation of parasitaemia with MSPDBL2 expression in different experimental replicate assays for different lab isolates. Spearman's r (rho) correlation coefficients and P values are shown for each isolate, the % parasitaemia/MSPDBL2 expression are shown in table S4.

| **Laboratory isolate** | **No. of assays** | **Correlation between parasitemia and MSPDBL2 expression in schizonts** | | **Correlation between parasitemia fold change and MSPDBL2 expression in schizonts** | |
|---|---|---|---|---|---|
| | | **rho** | **P** | **rho** | **P** |
| T9_96 | 5 | -0.05 | 1.00 | 0.1 | 0.95 |
| Palo Alto | 5 | 0.2 | 0.78 | -0.20 | 0.73 |
| GB4 | 7 | 0.32 | 0.51 | 0.42 | 0.35 |
| 7G8 | 5 | 0.36 | 0.56 | -0.67 | 0.26 |

**Table S7.** Effects of adding choline to serum-free medium on proportions of MSPDBL2+ schizonts in the HB3 and NF54 *P. falciparum* lines. Microscopical IFA counting of schizonts with at least 8 nuclei was performed on multiple experimental replicates of each line, with statistical analysis of proportions and effects within each replicate separately. The proportion of MSPDBL2+ schizonts are shown together with the 95% CI and are matched with associated gametocyte conversion rate (GCR) data measured in the same cultures as the proportion of stage 1 gametocytes developing in the first cycle of re-invasion using anti-Pfs16 staining (GCR data from Stewart et al. 2022 *Microbiology Spectrum* e0223422).

| Replicate Assays | Condition | MSPDBL2 +ve | DAPI +ve | Proportion of MSPDBL2+ schizonts (± 95%CI) | Fisher's Exact Test | Pfs16 +ve (GCR)* | Pfs16 -ve (GCR)* | Pfs16 GCR * | Fisher's Exact Test (p value) |
|---|---|---|---|---|---|---|---|---|---|
| | | HB3 | | | | | | | |
| R1 | - choline | 100 | 842 | 11.8 (9.8 – 14.2) | 0.0004 | 16 | 967 | 1.6 (1.0 – 2.6) | 0.141 |
| | + choline | 65 | 934 | 6.9 (5.5 – 8.8) | | 7 | 864 | 0.8 (0.4 – 1.6) | |
| R2 | - choline | 103 | 902 | 11.4 (9.5 -13.7) | < 0.00001 | 34 | 902 | 3.6 (2.6 – 5.0) | < 0.00001 |
| | + choline | 46 | 953 | 4.8 (3.6 – 6.4) | | 3 | 993 | 0.3 (0.1 – 0.9) | |
| R3 | - choline | 128 | 881 | 14.5 (12.3 – 17.0) | < 0.00001 | 36 | 929 | 3.7 (2.7 – 5.1) | 0.013 |
| | + choline | 63 | 976 | 6.4 (5.1 – 8.2) | | 18 | 950 | 1.9 (1.2 – 2.9) | |
| R4 | - choline | 172 | 1120 | 15.3 (13.3 – 17.6) | < 0.00001 | 24 | 954 | 2.5 (1.7 – 3.6) | 0.446 |
| | + choline | 63 | 930 | 6.8 (5.3 – 8.6) | | 19 | 961 | 1.9 (1.2 – 3.0) | |
| R5 | - choline | 34 | 758 | 4.5 (3.2 – 6.2) | 0.900 | 20 | 959 | 2.0 (1.3 – 3.1) | 0.8763 |
| | + choline | 32 | 739 | 4.3 (3.1 – 6.1) | | 21 | 957 | 2.1 (1.4 – 3.3) | |
| R6 | - choline | 60 | 939 | 6.4 (5.0 - 8.1) | 0.055 | 19 | 962 | 1.9 (1.2 – 3.0) | 0.0323 |
| | + choline | 81 | 925 | 8.7 (7.1 - 10.7) | | 8 | 983 | 0.8 (0.4 – 1.6) | |
| R7 | - choline | 76 | 923 | 8.2 (6.6 – 10.1) | 0.223 | 29 | 943 | 3.0 (2.1 – 4.3) | 0.0004 |
| | +choline | 90 | 909 | 9.9 (8.1 – 12) | | 8 | 983 | 0.8 (0.4 – 1.6) | |
| **Mean** | **– choline** | | | **12** | | | | **2.6** | |
| | **+ choline** | | | **7.9** | | | | **1.2** | |

| Replicate Assays | Condition | MSPDBL2 +ve | DAPI +ve | Proportion of PfMSPDBL2+ schizonts (± 95%CI) | Fisher's Exact Test | Pfs16 +ve (GCR)* | Pfs16 -ve (GCR)* | Pfs16 GCR* | Fisher's Exact Test (p value) |
|---|---|---|---|---|---|---|---|---|---|
| | | NF54 | | | | | | | |
| R1 | - choline | 2 | 577 | 0.3 (0.0 – 1.3) | 0.062 | 27 | 945 | 2.8 (1.9 – 4.0) | 0.0083 |
| | + choline | 13 | 962 | 1.3 (0.7 - 2.3) | | 11 | 978 | 1.1 (0.6 – 2.0) | |
| R2 | - choline | 2 | 997 | 0.2 (0.0 – 0.8) | 0.272 | 30 | 943 | 3.1 (2.2 – 4.4) | 0 |
| | + choline | 2 | 343 | 0.6 (0.0 – 2.2) | | 6 | 990 | 0.6 (0.3 – 1.3) | |
| R3 | - choline | 2 | 776 | 0.2 (0.0 – 1.0) | 1.0 | 40 | 819 | 4.7 (3.4 – 6.3) | 0 |
| | + choline | 3 | 996 | 0.3 (0.0 – 0.9) | | 13 | 973 | 1.3 (0.8 – 2.2) | |
| R4 | - choline | 6 | 984 | 0.6 (0.2 – 1.3) | 0.178 | 43 | 913 | 4.5 (3.4 – 6.0) | < 0.00001 |
| | + choline | 2 | 988 | 0.2 (0.0 – 0.8) | | 8 | 983 | 0.8 (0.4 – 1.6) | |
| R5 | - choline | 9 | 973 | 0.9 (0.4 – 1.8) | 0.628 | 70 | 861 | 7.5 (6.0 – 9.4) | 0.0219 |
| | + choline | 4 | 973 | 0.4 (0.1 - 1.1) | | 47 | 905 | 4.9 (3.7 – 6.5) | |
| R6 | - choline | 0 | 863 | 0.0 (0.0 – 0.5) | N/A | 8 | 983 | 0.8 (0.4-1.6) | 0.5798 |
| | + choline | 0 | 640 | 0.0 (0.0 – 0.7) | | 5 | 986 | 0.5 (0.2 – 1.2) | |
| **Mean** | **- choline** | | | **0.4** | | | | **3.9** | |
| | **+ choline** | | | **0.5** | | | | **1.5** | |

**Table S8.** Immunofluorescence microscopy counts of proportions of MSPDBL2 +ve schizonts in genetically-engineered parasite lines (among all those with at least 8 nuclei), using α-MSPDBL2 antibodies raised against conserved sequence N-terminal (Nter) or C-terminal (Cter) MSPDBL2 recombinant proteins. The tested lines are: the MSPDBL2 tagged line (MSPDBL2-TAG) and the MSPDBL2 disrupted line (MSPDBL2 DEL). Replicates and clones are indicated in brackets.

| Assay number | Line (and antibody) | MSPDBL2 +ve | DAPI +ve | Proportion of MSPDBL2+ schizonts (95% CI) |
|---|---|---|---|---|
| **1** | MSPDBL2-TAG rep1 (Nter) | 331 | 508 | 65.2 (60.9 – 69.2) |
| | MSPDBL2-TAG rep1 (Cter) | 232 | 368 | 63.0 (58.0 – 67.8) |
| | MSPDBL2-TAG rep2 (Nter) | 233 | 341 | 68.3 (63.2 – 73.0) |
| | MSPDBL2-TAG rep2 (Cter) | 145 | 183 | 79.2 (72.8 – 84.5) |
| | MSPDBL2 DEL Cl1 (Nter) | 480 | 520 | 92.3 (89.7-94.3) |
| | MSPDBL2 DEL Cl1 (Cter) | 0 | 930 | 0.0 |
| | MSPDBL2 DEL Cl3 (Nter) | 373 | 403 | 92.6 (89.5-94.7) |
| | MSPDBL2 DEL Cl3 (Cter) | 0 | 570 | 0.0 |
| **2** | MSPDBL2-TAG rep1 (Nter) | 261 | 309 | 84.5 (80.0-88.1) |
| | MSPDBL2-TAG rep1 (Cter) | 246 | 314 | 78.3 (73.4-82.6) |
| | MSPDBL2 DEL Cl1 (Nter) | 448 | 551 | 81.3 (77.8-84.3) |
| | MSPDBL2 DEL Cl1 (Cter) | 0 | 640 | 0.0 |
| **Mean** | **MSPDBL2-TAG (Nter)** | | | **72.6** |
| | **MSPDBL2-TAG (Cter)** | | | **73.5** |
| | **MSPDBL2 DEL (Nter)** | | | **88.7** |
| | **MSPDBL2 DEL (Cter)** | | | **0** |

**Table S9.** Gametocyte conversion rates of genetically engineered lines with multiple experimental replicates in the presence and absence of Shield-1 reagent. The tested lines are: 3D7/iGP (parental line), the MSPDBL2 tagged line (MSPDBL2-TAG) and the MSPDBL2 disrupted line (MSPDBL2 DEL).

| Assay number | Line (and treatment) | Pfs16 +ve | DAPI +ve | Pfs16 GCR (95% CI) |
|---|---|---|---|---|
| 1 | 3D7/iGP rep1 (- Shield-1) | 47 | 992 | 4.7 (3.6 –6.2) |
| | 3D7/iGP rep1 (+ Shield-1) | 349 | 690 | 50.6 (46.9 – 54.3) |
| | MSPDBL2-TAG rep1 (- Shield-1) | 104 | 791 | 13.1 (11.0 – 15.7) |
| | MSPDBL2-TAG rep1 (+ Shield-1) | 310 | 676 | 45.9 (42.1 –49.6) |
| | MSPDBL2 DEL Cl1 (- Shield-1) | 62 | 937 | 6.6 (5.2-8.4) |
| | MSPDBL2 DEL Cl1 (+ Shield-1) | 349 | 670 | 52.1 (48.3-55.8) |
| 2 | 3D7/iGP rep 2 (- Shield-1) | 39 | 929 | 4.2 (3.1 – 5.7) |
| | 3D7/iGP rep 2 (+ Shield-1) | 302 | 507 | 59.6 (55.2-63.7) |
| | MSPDBL2-TAG rep2 (- Shield-1) | 81 | 806 | 10.0 (8.1–12.3) |
| | MSPDBL2-TAG rep2 (+ Shield-1) | 402 | 837 | 48.0 (44.7–51.4) |
| | MSPDBL2 DEL Cl2 (- Shield-1) | 88 | 913 | 9.6 (7.9-11.7) |
| | MSPDBL2 DEL Cl2 (+ Shield-1) | 289 | 670 | 43.1 (39.4-46.9) |
| 3 | 3D7/iGP rep 3 (- Shield-1) | 93 | 849 | 10.9 (9.0-13.2) |
| | 3D7/iGP rep 3 (+ Shield-1) | 193 | 568 | 34.0 (30.2-38.0) |
| | MSPDBL2-TAG rep3 (- Shield-1) | 73 | 903 | 8.1 (6.5-10.0) |
| | MSPDBL2-TAG rep3 (+ Shield-1) | 269 | 758 | 35.5 (32.2-39.0) |
| | MSPDBL2 DEL Cl3 (- Shield-1) | 66 | 901 | 7.3 (5.8-9.2) |
| | MSPDBL2 DEL Cl3 (+ Shield-1) | 249 | 775 | 32.13 (29.0-35.5) |
| 4 | 3D7/iGP rep1 (- Shield-1) | 61 | 938 | 6.5 (5.1-8.3) |
| | 3D7/iGP rep1 (+ Shield-1) | 367 | 662 | 55.4 (51.6-59.2) |
| | MSPDBL2-TAG rep1 (- Shield-1) | 43 | 956 | 4.5 (3.3 - 6.0) |
| | MSPDBL2-TAG rep1 (+ Shield-1) | 358 | 651 | 55.0 (51.1 - 58.8) |
| | MSPDBL2 DEL Cl1 (- Shield-1) | 60 | 942 | 6.4 (5.0 - 8.1) |
| | MSPDBL2 DEL Cl1 (+ Shield-1) | 330 | 626 | 52.7 (48.8 - 56.6) |
| 5 | 3D7/iGP rep 2 (- Shield-1) | 77 | 931 | 8.3 (6.7-10.2) |
| | 3D7/iGP rep 2 (+ Shield-1) | 383 | 643 | 59.6 (55.7-63.3) |
| | MSPDBL2-TAG rep2 (- Shield-1) | 66 | 935 | 7.1 (5.6- 8.9) |
| | MSPDBL2-TAG rep2 (+ Shield-1) | 366 | 639 | 57.3 (53.4-61.1) |
| | MSPDBL2 DEL Cl2 (- Shield-1) | 44 | 583 | 7.5 (5.6 - 10.0) |
| | MSPDBL2 DEL Cl2 (+ Shield-1) | 5 | 12 | 41.7 (19.3 -68.1) |
| 6 | 3D7/iGP rep 3 (- Shield-1) | 80 | 900 | 8.9 (7.2-10.9) |
| | 3D7/iGP rep 3 (+ Shield-1) | 336 | 703 | 47.8 (44.1 - 51.5) |
| | MSPDBL2-TAG rep3 (- Shield-1) | 47 | 951 | 5.0 (3.7 - 6.5) |
| | MSPDBL2-TAG rep3 (+ Shield-1) | 315 | 627 | 50.2 (46.3 - 54.1) |
| | MSPDBL2 DEL Cl3 (- Shield-1) | 70 | 875 | 8.0 (6.4 - 10.0) |
| | MSPDBL2 DEL Cl3 (+ Shield-1) | 365 | 641 | 56.9 (53.1- 60.7) |
| **Mean** | **3D7/iGP (- Shield-1)** | | | **7.2** |
| | **3D7/iGP (+ Shield-1)** | | | **51.1** |
| | **MSPDBL2-TAG (- Shield-1)** | | | **7.9** |
| | **MSPDBL2-TAG (+ Shield-1)** | | | **48.6** |
| | **MSPDBL2 DEL (- Shield-1)** | | | **7.6** |
| | **MSPDBL2 DEL (+ Shield-1)** | | | **47.4** |

**Table S10.** Multiplication rates of genetically-engineered *P. falciparum* lines in 6-day assays of exponential growth. Parasite genome copy numbers obtained by qPCR analysis of the single-copy gene PF3D7_1031000 (*Pfs25*) are presented. Multiplication rates were calculated by logistic regression using data from all three replicates for each line.

| | Genome copy numbers in 1µl DNA from each of three replicate growth assays | | | | |
|---|---|---|---|---|---|
| **Parasite Line** | **Log qPCR Day 0** | **Log qPCR Day 2** | **Log qPCR Day 4** | **Log qPCR Day 6** | **Multiplication rate (95% CI) per 48h** |
| 3D7 | 2.71 | 3.96 | 4.66 | 5.31 | 7.26 (5.89-8.96) |
| | 2.87 | 4.00 | 4.65 | 5.65 | |
| | 2.85 | 3.96 | 4.75 | 5.37 | |
| 3D7/iGP | 1.86 | 2.84 | 3.26 | 4.15 | 6.53 (5.22-8.17) |
| | 1.65 | 2.73 | 3.67 | 4.52 | |
| | 1.89 | 2.87 | 3.36 | 4.28 | |
| MSPDBL2-TAG (rep1) | 2.37 | 3.79 | 4.70 | 5.16 | 8.81 (6.61-11.76) |
| | 2.45 | 3.59 | 4.82 | 5.33 | |
| | 2.57 | 3.69 | 5.33 | 5.28 | |
| MSPDBL2-TAG (rep2) | 2.83 | 3.59 | 4.60 | 5.21 | 7.91 (6.24-10.02) |
| | 2.60 | 3.91 | 4.89 | 5.62 | |
| | 2.67 | 3.74 | 4.56 | 5.28 | |
| MSPDBL2-TAG (rep3) | 2.50 | 3.73 | 4.43 | 4.99 | 7.46 (6.14-9.08) |
| | 2.57 | 3.63 | 4.50 | 5.25 | |
| | 2.51 | 3.65 | 4.48 | 5.28 | |
| MSPDBL2 DEL (clone 1) | 2.24 | 3.22 | 4.05 | 4.86 | 8.82 (6.70-11.60) |
| | 2.02 | 3.33 | 4.33 | 5.08 | |
| | 2.27 | 2.64 | 4.14 | 4.94 | |
| MSPDBL2 DEL (clone 2) | 2.14 | 3.23 | 3.80 | 4.86 | 6.48 (5.19-8.08) |
| | 2.12 | 3.29 | 4.18 | 4.98 | |
| | 2.22 | 3.32 | 3.85 | 4.93 | |
| MSPDBL2 DEL (clone 3) | 2.27 | 3.29 | 4.12 | 4.56 | 7.84 (6.46-9.50) |
| | 2.43 | 3.45 | 4.16 | 4.97 | |
| | 2.54 | 3.37 | 4.35 | 4.99 | |

**Table S11.** Sequences of primers designed for cloning and genotyping in the present study.

| name | Primers | | | experiment | |
|---|---|---|---|---|---|
| | sequence | orientation | restriction sites | | |
| | | | | Generation of transgenic parasites | |
| | | | | construct/use | vector |
| P1 | TGACTCCAAGCAGAGGTACCCTTGTACAGCTCGTCC | F | BamHI | pL7M2_TdTom_T2A/G418 | pL7M2_T2A/G418 |
| P2 | TATTCTTTTAGGATCCGTGAGCAAGGGCGA | R | KpnI | pL7M2_TdTom_T2A/G418 | pL7M2_T2A/G418 |
| P3 | CCGCAGATCTCTCGAGAAAACATTATCTCCTTCTGAA | F | XhoI | *mspdbl2* -pL7M2_TdTom_T2A/G418 | pL7M2_TdTom_T2A/G418 |
| P4 | CCTTGCTCACGGATCcATTTTTAAATAAATTTGTAATATCTTTTGAT | R | BamHI | *mspdbl2* -pL7M2_TdTom_T2A/G418 | pL7M2_TdTom_T2A/G418 |
| P5 | GGACTGAACAATCTATTAAATATAACAATG | F | | | |
| P6 | GTATTCAATTATGCTTTATAAGAAACACA | R | | genotype of *mspdbl2 -TdTom-3xHA* (T2A_G418) | |
| P7 | ATGTTGTTGTCCTCGGAGGAG | R | | | |
| P8 | TATTCTTTTAGGATCCAATGATATATATTTTATCTATTGTATTTTATATATTTTTTT | F | BamHI | *mspdbl2_DEL* -pL7M2_T2A/G418 | pL7M2_T2A/G418 |
| P9 | TGACTCCAAGCAGAGGTACCACAAATCTCTTCAGTAGG | R | KpnI | *mspdbl2_DEL* -pL7M2_T2A/G418 | pL7M2_T2A/G418 |
| P10 | GTTCATTGAACTGTTAATGTGAATGTTTTTG | F | | | |
| P11 | CTTGTTCAATCATTGGTCCTGGA | R | | genotype of *mspdbl2 del* (T2A_G418) | |
| P12 | CGTACGAAAATCTTGTAATG | R | | | |

**Table S12.** *mspdbl2-tag* and *mspdbl2 del* sequences inserted into the endogenous locus:

*A. mspdbl2-TdTom-3xHA* sequence (3867 bp)*:*

ATGATATATATTTTATCTATTGTATTTTATATATTTTTTTTACATATTGATATATATGTAAACATTTATTCTACATGTTTTGTTGTAAATGAGGGGAACCCTAATTTA
AGAAATAACATAATTAATGATGATGAACTAAAGGGGAAAGCATATAATAATACTATAGATGCTAATAACCAAAATATAGAATATAATAAAAACTTAAAGCACA
ATGTAAACTCATCTCATATATCTAAATTTTCGGATATTATGGATCAAGAAGATAAAGGAGATAATGAAAATTCTCATGACATAAAATTTGAAGAAAAAAAAAAT
ATTAATAAATCTTTAGACGCTGAATCCAATTATGGTATTAATGAAATTAGTATTACTGGTAATGATAGTAATAGTGATAATAGTAATCAGAATATTTTTCCAGAT
GGTAGTGAATTAGCTGGAGGTATTCCTCGTTCTATATATACTATTAACCTTGGTTTTAATAAATGTCCTACTGAAGAGATTTGTAAAGACTTTAGTAATCTTCCAC
AATGTCGAAAGAATGTACATGAAAGAAATAATTGGTTGGGCTCAAGTGTAAAAAATTTTTCAAGTGATAATAAGGGGGTTCTTGTTCCTCCAAGAAGACAATCT
TTATGTTTAAGAATTACATTACAAGATTTTCGTACGAAAAAGAAAAAGGAAGGAGATTTTGAAAAATTTATTTATTCATATGCATCATCTGAAGCTAGAAAATTA
AGAACCATACACAATAATAACTTAGAAAAAGCTCATCAAGCTATAAGATATAGTTTTGCAGATATTGGAAATATTATTAGAGGAGATGACATGATGGATACACC
TACGTCAAAAGAAACCATAACATATTTAGAAAAAGTACTTAAAATTTATAATGAAAATAATGATAAACCAAAAGATGCAAAAAAAATGGTGGACAGAAAACAGG
CATCATGTTTGGGAAGCAATGATGTGCGGATATCAGAGTGCGCAGAAAGATAACCAATGTACAGGTTATGGTAACATTGATGATATACCACAATTTTTAAGGT
GGTTCAGAGAGTGGGGAACATATGTCTGTGAAGAAAGCGAAAAAAAATATGAACACACTAAAAGCTGTTTGCTTTCCGAAACAGCCAAGAACCGAAGCGAATC
CTGCATTGACTGTACATGAAAATGAAATGTGCTCATCAACTTTAAAAAAAATATGAAGAATGGTATAATAAAAGGAAAACTGAATGGACTGAACAATCTATTAAA
TATAACAATGACAAAATTAATTATACAGATATAAAAACATTATCTCCTTCTGAATATTTAATAGAAAAATGTCCTGAATGTAAATGTACCAAAAAAAAATTTGCAA
GATGTATTTGAACTTACATTTGATGGAAAAGCTTTATTAGAAAAGCTAAAAAAAGAAGAATCACCTGTGAGTAATAGTGTGAATGCCTTACCTGAACCAGGTCA
AATTACATTACCTGATCCTTCATTAAAACAAACAACACAACAGGAAAATCAACCTGTTGTAGAAACACCTGTTACCACAGCTGTTATTAATGAACATCAAGGACA
AACAGAACCGAATAAAGGTGACAACAATAATGAAAGAGAAAATCATGAAAGTAATGTTGGTAGCATCCAAGAAGTAAACCAAGGTAGCGTGAGCGAAGAAT
CACATTCTAAAACTATAGATCCTTCTAAGATTGACGACCGTTTGGAATTAAGTAGTGGGTCATCATCTCTTGAACAACACTCTAAGGAAGATGTAAAAAAGGGA
TGTGCTTTAGAATTGGTACCTTTATCTTTATCGGATATTGAACAGATAGCTAATGAAAGCGAAGATGTACTGGAAGAGATAGAAGAAGAAATTAATACAGATG
GGGAAATAGAATATATAACAGAAGAAGAAATAAAAGAAGATATAGAAGAAGAAACAGAAGAAGATATAGAAGAAGAAACAGAAGAAGAAACAGAAGAAGA
AACAGAAGAAGAAGCAGATGAAGAAACAGTAAAAGAAATAGAAGACAAACCAGAACAAGAAATTAAAAATAAATCGCTAGAAGAAAAACAAATAGATAAAA
ATACAGATACCAGTGAAAAGAAAGGATTTAATAATTCAGAAAAAGATGAAAAAGCTCGAAATTTAATTTCTAAAAATTATAAAAATTATAATGAACTAGATAAA
AACGTTCATACTTTAGTAAATTCAATTATTAGTTTATTAGAAGAAGGTAATGGAAGTGATTCTACCTTGAATAGTTTATCAAAAGATATTACAAATTTATTTAAAA
ATGGATCCGTGAGCAAGGGCGAGGAGGTCATCAAAGAGTTCATGCGCTTCAAGGTGCGCATGGAGGGCTCCATGAACGGCCACGAGTTCGAGATCGAGGGC
GAGGGCGAGGGCCGCCCCTACGAGGGCACCCAGACCGCCAAGCTGAAGGTGACCAAGGGCGGCCCCCTGCCCTTCGCCTGGGACATCCTGTCCCCCCAGTTC
ATGTACGGCTCCAAGGCGTACGTGAAGCACCCCGCCGACATCCCCGATTACAAGAAGCTGTCCTTCCCCGAGGGCTTCAAGTGGGAGCGCGTGATGAACTTCG
AGGACGGCGGTCTGGTGACCGTGACCCAGGACTCCTCCCTGCAGGACGGCACGCTGATCTACAAGGTGAAGATGCGCGGCACCAACTTCCCCCCCGACGGCC
CCGTAATGCAGAAGAAGACCATGGGCTGGGAGGCCTCCACCGAGCGCCTGTACCCCCGCGACGGCGTGCTGAAGGGCGAGATCCACCAGGCCCTGAAGCTG
AAGGACGGCGgccactacctggtggagttcaagACCATCTACATGGCCAAGAAGCCCGTGCAACTGCCCGGCTACTACTACGTGGACACCAAGCTGGACATCACCTCC
CACAACGAGGACTACACCATCGTGGAACAGTACGAGCGCTCCGAGGGCCGCCACCACCTGTTCCTGGGGCATGGCACCGGCAGCACCGGCAGCGGCAGCTCC
GGCACCGCCTCCTCCGAGGACAACAACATGGCCGTCATAAAAGAATTTATGAGATTCAAGGTACGTATGGAAGGTTCTATGAATGGACATGAGTTTGAGATTG
AAGGAGAAGGTGAAGGTCGTCCTTATGAGGGAACCCAAACTGCAAAACTTAAGGTAACTAAAGGTGGTCCATTGCCATTTGCTTGGGATATATTGAGTCCACA
ATTCATGTACGGTAGTAAGGCCTACGTAAAACATCCAGCCGATATTCCAGATTACAAAAAGCTTTCATTCCCTGAGGGATTCAAGTGGGAAAGAGTTATGAATT
TCGAAGATGGTGGTCTTGTAACCGTAACCCAAGATTCTTCACTTCAAGATGGAACTTTAATATATAAAGTAAAAATGAGAGGAACCAATTTCCCTCCTGATGGA
CCTGTAATGCAAAAAAAAACAATGGGTTGGGAGGCAAGTACAGAGAGATTGTACCCACGTGATGGTGTTCTTAAGGGAGAGATCCATCAAGCTTTGAAATTG
AAGGATGGTGGACATTACTTGGTAGAGTTTAAGACTATATATATGGCAAAGAAGCCAGTTCAATTGCCTGGTTATTACTACGTTGATACTAAGTTAGATATTAC
AAGTCATAATGAGGATTATACCATAGTAGAACAATATGAACGTAGTGAAGGAAGACACCATTTGTTTCTGTACGGCATGGACGAGCTGTACAAGggtaccTCTGC
TTGGAGTCATCCTCAATTTGAAAAAGGAGGATCTAGTTACCCTTACGATGTTCCTGACTATGCGGGCTATCCCTATGACGTCCCGGACTATGCCATGGGCTACCC
TTACGACGTTCCAGATTACGCTGGAGGTTCTGGT

Sequence inserted into pL2M2_TdTom_T2A/G418 (BamH1/Td-TOM/Kpn1/3xHA tag)

Predicted MW of translated product: 147.2 KDa

*B. mspdbl2 del* sequence *(681 bp):*

ATGATATATATTTTATCTATTGTATTTTATATATTTTTTTTACATATTGATATATATGTAAACATTTATTCTACATGTTTTGTTGTAAATGAGGGGAACCCTAATTTA
AGAAATAACATAATTAATGATGATGAACTAAAGGGGAAAGCATATAATAATACTATAGATGCTAATAACCAAAATATAGAATATAATAAAAACTTAAAGCACA
ATGTAAACTCATCTCATATATCTAAATTTTCGGATATTATGGATCAAGAAGATAAAGGAGATAATGAAAATTCTCATGACATAAAATTTGAAGAAAAAAAAAAT
ATTAATAAATCTTTAGACGCTGAATCCAATTATGGTATTAATGAAATTAGTATTACTGGTAATGATAGTAATAGTGATAATAGTAATCAGAATATTTTTCCAGAT
GGTAGTGAATTAGCTGGAGGTATTCCTCGTTCTATATATACTATTAACCTTGGTTTTAATAAATGTCCTACTGAAGAGATTTGTggtaccTCTGCTTGGAGTCATCC
TCAATTTGAAAAAGGAGGATCTAGTTACCCTTACGATGTTCCTGACTATGCGGGCTATCCCTATGACGTCCCGGACTATGCCATGGGCTACCCTTACGACGTTCC
AGATTACGCTGGAGGTTCTGGT

*mspdbl2* sequence inserted into pL2M2_T2A/G418 (Kpn1/3xHA tag)

Predicted MW of translated product: 24.5 KDa
